# Osmolar modulation drives reversible cell cycle exit and human pluripotent cell differentiation via NF-κВ and WNT signaling

**DOI:** 10.1101/2023.04.14.536882

**Authors:** Jonathan Sai-Hong Chui, Teresa Izuel-Idoype, Alessandra Qualizza, Rita Pires de Almeida, Bernard K. van der Veer, Gert Vanmarcke, Paraskevi Athanasouli, Ruben Boon, Joris Vriens, Kian Peng Koh, Leo van Grunsven, Catherine M. Verfaillie, Frederic Lluis

## Abstract

Terminally differentiated cells are regarded as the most stable and common cell state in adult organisms as they reside in growth arrest and carry out their cellular function. Improving our understanding of the mechanisms involved in promoting cell cycle exit would facilitate our ability to manipulate pluripotent cells into mature tissues for both pharmacological and therapeutic use. Here, we demonstrated that a hyperosmolar environment enforced a protective p53-independent quiescent state in dedifferentiated hepatoma cells and pluripotent stem cells (PSCs)-derived models of human hepatocytes and endothelial cells, representing the endodermal and mesodermal lineages. Prolonged culture in hyperosmolar conditions stimulated transcriptional and functional cell maturation. Interestingly, hyperosmolar conditions did not only trigger cell cycle exit and cellular maturation but were also necessary to maintain this maturated state, as switching back to plasma osmolarity caused the loss of maturation markers and the gain of proliferative markers. Transcriptome analysis revealed activation of NF-κВ and repression of WNT signaling as the two main pathways downstream of osmolarity-regulated growth arrest and cell maturation, respectively. This study revealed that increased osmolarity serves as a biochemical signal to promote long-term growth arrest, transcriptional changes, and maturation into different lineages, serving as a practical method to generate differentiated hiPSCs that resemble their mature counterpart more closely.

## INTRODUCTION

During embryonic development and organogenesis, differentiating cells exit the cell cycle and enter a prolonged or permanent G_0_ growth arrest to achieve their final mature state, which represents the most common cell state in adult organs^1, 2^. The transition from proliferation to cell cycle exit is accompanied by a transcriptional and functional switch in essential regulators of the cell cycle program. To exit the cell cycle and to maintain a post-mitotic state, the pro-mitotic cell cycle genes such as cyclins and cyclin-dependent kinases (CDKs) are suppressed, while cell cycle inhibitors like CDK inhibitors (CDKi) become upregulated. These bind to and inhibit the activity of cyclin/CDK complexes and negatively regulate cell cycle progression. Not only are CDKi essential in regulating proliferation during development, but they are also key players in promoting differentiation, cell quiescence, senescence, and inhibiting apoptosis^3^. Among the CDKi, the INK4 (p16^INK4A^, p15^INK4B^, p18^INK4C^ and p19^INK4D^) and the CIP/KIP (p21^CIP^1^/WAF1^, p27^KIP1^ and p57^KIP2^) families have been described to regulate cell cycle progression and cell cycle exit. Following exit from the cell cycle, the growth arrest of differentiated cells can further transgress into a reversible (quiescent) or irreversible (senescent) state^4–7^. Yet, it remains unclear how this growth arrest is initiated and how cells in mature tissues maintain a stable non-proliferative state.

The ability to cultivate and differentiate human induced pluripotent stem cells (hiPSC) towards all three embryonic lineages (endoderm, mesoderm, and ectoderm) has provided a fundamental base to generate tissue-specific cells not only to model diseases but also for regenerative and drug screening purposes^8^. Despite their great potential, current differentiation conditions generate cells typically displaying an immature resemblance to their *in vivo* counterparts expressing unsatisfying levels of maturation and growth arrest markers. Although hiPSC differentiation protocols depend on a consecutive modulation of signaling pathways by the addition of growth factors^9–11^, several reports demonstrated that environmental factors, such as oxygen levels and surface stiffness, can greatly impact stem cell self-renewal and differentiation^12, 13^. However, the effects of other culturing factors on pluripotent cell differentiation, such as medium osmolarity, have not yet been explored.

Osmolarity is a measure for the pressure of a solution defined by its diffusion across the cell membrane. Generally, hiPSC differentiation protocols are performed in media with osmolar values resembling plasma levels (285-300 mOsm)^14^. In contrast, cells residing in adult tissues have developed molecular mechanisms to adapt to osmolar fluctuations, and several studies have reported that the interstitial fluid within multiple adult organs (liver, kidney, cornea, gastrointestinal tract, intervertebral discs, among others) has significantly higher osmolar levels due to their metabolic activity^15–19^. As tissue-resident cells reside in such hyperosmolar (HypOsm) conditions, this may indicate that hyperosmolarity might be a crucial factor to be considered for obtaining and maintaining more mature cultured cells for prolonged periods of time.

As the central metabolic hub, the liver acts as an active buffer of systemic changes by its ability to modify glucose levels, detoxify xenobiotics, facilitate digestion, and to synthesize proteins crucial for excretion^20^. Therefore, the liver is found to be within the hyperosmolar range to facilitate the exchange of metabolites and proteins between the hepatic environment and the plasma under healthy conditions^18, 20^. As current liver models fail to properly recapitulate the human liver, achieving improved liver models is of high interest to serve as tools in disease modeling or drug toxicity screenings.

Despite the high osmolar environments in various physiological tissues, hyperosmolarity is *in vitro* associated with cellular stress and has been previously described drive mammalian cells to undergo cell shrinkage, intracellular dehydration, oxidative stress, DNA damage, and caspase activation^21^. As cellular damage accumulates, increased p53 levels promote growth arrest, and cells become primed to undergo apoptosis^22^. However, other studies have stated that under the right conditions mature mammalian cells are able to overcome osmolar stress and prevent apoptosis by eliciting osmo-adaptive responses^23–25^. These adaptative strategies are mainly orchestrated by the NFAT5 transcription factor, also known as the tonicity-responsive enhancer-binding protein (TonEBP)^25, 26^. NFAT5 targets tonicity-responsive gene transcription and regulates the expression of osmo-protective genes that are responsible for raising intracellular levels of osmolytes in the cell, preventing cell shrinkage, and allowing osmo-adaptation and cellular survival^25, 27^. As hyperosmolarity drives cells to either adapt or perish^28^, this indicates that the in vitro cellular response to HypOsm conditions could be cell type or dosage dependent and should be titrated as such for each case.

In this study, we describe the use of HypOsm culture conditions to promote the maturation of hepatoma and hiPSC-derived differentiation driven by a shared underlying mechanism. We first show in hepatoma lines and hiPSC differentiation to hepatocyte-like cells (HLC) that in contrast to toxic hyperosmolarity, the prolonged culture within physiological osmolarity levels stabilizes a G_0_ growth arrest and hepatic transcriptional maturation in a p53-independent manner without affecting cell viability. Next, we discovered WNT and NF-κВ signaling as the two main downstream effectors responsible for these cellular changes. Interestingly, physiological HypOsm conditions are necessary for both the initiation and the stabilization of the growth arrest and enhanced maturation in hiPSC differentiation protocols. Finally, we demonstrate that physiological hyperosmolarity also improves pluripotent to endothelial cell maturation. Altogether, we show that instead of being considered a stress signaling, physiological HypOsm levels promote a comprehensive transcriptional re-wiring that is beneficial for inducing hepatic and endothelial PSC growth arrest and differentiation.

## METHODS

### Hepatoma cell line culture

HUH7.5 and Hep3B lines were plated at a seeding density of 10.4 × 10^4^ cells/cm^2^ or 50.000 cells/mL in DMEM medium low glucose (Gibco, 31885023) supplemented with 10% FBS and 1X penicillin-streptomycin. Cell lines were passaged following dissociation using 0.25% Trypsin-EDTA solution (Gibco, 15050065) upon reaching 70% confluency.

### Stem cell culture and passaging

The hiPSC line, Sigma 0028, was purchased from Sigma-Aldrich and maintained on coated plates with Corning™ Matrigel™ hESC-Qualified Matrix (Corning, 734-1440) in Essential 8™ Flex Medium (E8 Flex; Gibco™, A2858501). The cells were passaged at 65% confluency using 0.1% Ethylenediaminetetraacetic acid (EDTA, Life Technologies, 15575020) in PBS. The hiPSCs were genetically engineered to induce the overexpression of 3 transcription factors (TFs), namely *HNF1A*, *FOXA3*, and *PROX1* (named HC3X-PSC) as described earlier to generate hepatocyte-like progeny; or one TF, namely *ETV2* (named iETV2-PSC) to generate endothelial cells^29, 30^. When the cells reached 65% confluency, cells were dissociated into single cells using StemPro Accutase (Gibco, A2644501) cell dissociation reagent. Cells were plated on Matrigel Based Membrane Growth Factor Reduced coated plates (Corning, 734-0270) at a density of ±8.75 × 10^4^ cells/cm^2^ or 3 × 10^5^ cells/ml in mTeSR medium (Stem Cell Technologies, 85850) supplemented with 1:100 RevitaCell Supplement (Gibco, A2644501). When the cells obtained a 70% - 80% confluency, the cells were differentiated towards either the hepatocyte or endothelial lineage. The use of hiPSCs for research was approved by the “Commissie Medische Ethiek,” UZ KU Leuven/Onderzoek U.Z. Gasthuisberg, Herestraat 49, B 3000 Leuven, under file number S52426.

### Generation of FUCCI- and 7TGP-HUH7.5 reporter cell lines

For the generation of the transduced FUCCI (Addgene, Plasmid #86849) and 7TGP (Addgene, Plasmid #24305) HUH7.5 lines, we initially transfected HEK293T cells using the Fugene HD Transfection Reagent (Promega, E2311) according to the manufacturer’s recommendations. Donor plasmids were packaged with 2^nd^ generation lentiviral packaging vectors p-CMV-VSV-G (Addgene, Plasmid #8454) and pCMV-dR8.2 (Addgene, Plasmid #12263). Two days after transfection, the conditioned media of transfected HEK293T cells was filtered and transferred to HUH7.5 cells. Following puromycin selection, these lines were validated by flow cytometry. All plasmids are accessible on Addgene.

### hiPSC differentiation to the hepatocyte lineage

The differentiation was started using previously described cytokine regimes and was stopped after 30 days of differentiation^31^. Briefly, mTeSR medium was replaced by Liver Differentiation Medium (LDM), and 50 ng/mL WNT-3A (Peprotech, 315-20) and 50 ng/mL Activin A (Peprotech, 120-14E) was supplemented at day 0, followed by only 50 ng/mL Activin-A at day 2. At day 4, 50 mg/mL BMP4 (Peprotech, 120-05ET) and 5 µg/mL doxycycline (Sigma-Aldrich, D9891) were added. From day 8 until day 12, 20 ng/mL acidic FGF (aFGF) (Peprotech, 120-05ET) was supplemented to the medium. From day 12 onwards, cells were cultured with 20 ng/mL HGF (Peprotech, 100-39) until the end of the differentiation. All cytokines were purchased from Peprotech. The differentiation medium was supplemented with 0.6% DMSO (Merck, D2650) during the first 12 days of the culture and with 2% DMSO from day 12 onward under PlasOsm conditions. Medium was changed every two days of the differentiation.

### hiPSC differentiation to the endothelial lineage

On day 0 of differentiation, mTeSR medium was replaced by LDM containing 5 μg/ml doxycycline (Sigma-Aldrich, D9891) and 10 ng/ml basic Fibroblast Growth Factor (bFGF, Peprotech, 100-18C). From day two onwards, 2% Fetal Bovine Serum (FBS) was added, and from day 4 onwards, 1% Endothelial Cell Growth Supplement (ECGS, R&D systems, 390599) was added to the LDM medium. Cells were passed every 4 days until day 12 to avoid overgrowth. Medium was changed every two days of the differentiation.

### Medium composition and osmolar treatment

Hepatoma cell lines HUH7.5 and Hep3B under HypOsm conditions were treated with DMEM supplemented with 250 mM of glycine (Osm^G^, Sigma-Aldrich, 1041691000) or 250 mM mannitol (Osm^M^, Sigma-Aldrich, M4125) 16 hours after plated.

Hepatocyte-like cell (HLC) differentiation: cells treated with Osm^G^ and Osm^M^ were cultured in LDM supplemented with 250 mM of glycine (Osm^G^) or 250 mM mannitol (Osm^M^) on day 12 of the protocol. hiPSC-HLC cells were cultured until day 30 of differentiation. Upon hyperosmolar treatment, 2% DMSO was removed from day 12 onwards.

Endothelial differentiation: cells treated with Osm^G^ and Osm^M^ were supplemented with 1x non-essential amino acids (50X) (NEAA, Gibco, 11140050) and 1x essential amino acids (100X) (EAA, Gibco, 11130051) at day 10 of differentiation and until the end of the protocol. On day 14, 250 mM of glycine (Osm^G^) or 250 mM mannitol (Osm^M^) was added to the LDM medium. Cells were cultured until day 26 of differentiation.

### Osmolar reversion

Hepatoma cell line HUH7.5 was cultured for 5 days in 550 mOsm conditions in Osm^G^ and Osm^M^ media. To revert to plasma osmolarity, the medium was changed back to regular DMEM supplemented with 10% FBS and cultured for an additional 5 days.

hiPSC-HLCs under hyperosmolar treatments were gradually reverted to plasma osmolarity at day 20 by reducing osmolarity by 20% every other day until reaching plasma osmolarity at day 30 of differentiation.

### Osmolar measurements

The osmolarity of the media was measured with a vapor pressure osmometer (5 Wescor 5500, Artisan Technology Group, 69905-6). The calibration of the instrument was performed using sodium chloride standards of 290 and 1000 mOsm/kg H_2_O. Specifically, a filter paper disc with a 6.5 mm diameter was saturated with 10 µL of the to-be-measured solution and placed into the instrument. Murine kidneys, spleen, liver, and blood were harvested from 10-week-old C57BL/6 mice (ECD M012/2023). Collected blood was captured in heparin-coated Microvette tubes (Sarstedt, MV-E-500) and centrifuged for 10 minutes at 2000 × g. Harvested organs were mechanically homogenized and osmolarity levels were measured by soaking the filter paper disc in the homogenate.

### Immunofluorescence staining

Cells were fixed with 4% PFA for 15 minutes at room temperature. Samples were permeabilized and blocked in a single step with 0.2% Triton-X and 5% donkey serum (Jackson Immunoresearch, 017-000-121). Primary antibodies were subsequently added and incubated overnight at 4 °C. Next, cells were washed with PBS, followed by a 40 minute incubation with the secondary antibodies. After 10 minutes of DAPI exposure, cells were imaged using the High Throughput Imager Operetta CLS system (Perkin Elmer, HH16000000). Image analysis and quantification were performed using the Harmony 4.2 software (Perkin Elmer, HH17000001).

### Flow cytometry experiments

For Ki67 detection, cells were collected and 1 × 10^6^ cells per condition were fixed with 70% ethanol. Samples were washed with PBS with 2% FBS and blocked with 5% donkey serum at room temperature for 30 – 60 minutes. Cells were centrifuged at 1400 rpm for 4 minutes and incubated with Ki-67 Recombinant Rabbit Monoclonal Antibody (SP6) (ThermoFisher, MA5-14520) at room temperature for 30 minutes. Cells were centrifuged at 1400 rpm for 4 minutes, resuspended with PBS 2% FBS and incubated with the secondary antibody at room temperature in the dark for 20 minutes. Then, cells were centrifuged, washed with PBS 2% FBS, and resuspended with DAPI. Samples were then examined by flow cytometry using the BD FACSCanto HTS flow cytometer and analyzed by the FlowJo software.

For annexin V apoptosis analysis, cells were collected, counted to obtain 1·× 10^6^ cells, and resuspended in annexin V binding buffer (BD Pharmigen, 51-66121E). Cells were then incubated with APC-conjugated Annexin V (Thermo-eBioscience, BMS306APC-100) at room temperature for 15 minutes. Subsequently, cells were centrifuged and resuspended in binding buffer containing DAPI.

Cell cycle changes were additionally assessed using the 5-ethynyl-2’-deoxyuridine (EdU) assay using the Click-iT™ EdU Alexa Fluor™ 647 Flow Cytometry Assay Kit (ThermoFisher, C10418). HUH7.5 cells were incubated at 37 °C for 2 hours with 10 µM EdU using the Click-iT EdU Assay Kit (Thermofisher, C10418) according to the manufacturer’s recommendations.

All samples from different experiments were examined by flow cytometry using the BD FACSCanto HTS flow cytometer and analyzed by the FlowJo software.

### Functional assessments

CYP450-dependent metabolization was determined by using the fluorometric probe BFC (Merck, B5057) as previously described^32^, for which metabolization was assessed for an incubation time of 4 hours. AFP, albumin, and urea secretion were determined with the AFP Human ELISA Kit (Invitrogen, EHAFP) (1:3000 dilution), Albumin ELISA Kit (Bethyl Laboratories, Inc., BET E80-129) (1:50 dilution), and QuantiChrom^TM^ Urea Assay Kit (BioAssay Systems, DIUR-500), respectively. Proteins were allowed to accumulate for 2 days before determining protein concentration, with negative controls specified according to the manufacturer’s guidelines. All protein levels were normalized for total cell number. For all functional readouts, cell number was determined with a NucleoCounter Automated Cell Counter (Chemometec, NC-200).

### Western blot

Cells were collected by centrifugation upon dissociation with 0.25% Trypsin-EDTA solution. Whole-cell lysates were prepared using RIPA cell lysis buffer (Sigma-Aldrich, R0278) containing 1:100 phosphatase inhibitor cocktail 2 (Sigma-Aldrich, P5726), phosphatase inhibitor cocktail 3 (Sigma-Aldrich, P0044), and protease inhibitor (Sigma-Aldrich, P8340). Samples were rotated for at least 30 minutes at 4 °C and spun at maximum speed for 10 minutes at 4 °C. The supernatants of the samples were collected, and protein concentration was determined by the Bradford assay (Bio-Rad, 23246). Equal amounts per sample were combined with Laemmli buffer, denatured for 5 minutes at 95 °C, and subjected to SDS/PAGE separation, followed by immunoblotting. The antibodies used in this study are listed in **Supplementary Table 1**. β-Actin was used as an internal loading control. Protein quantification was performed with ImageJ software from N=3 independent biological replicates.

### RNA extraction and quantitative reverse-transcription PCR

Total RNA was extracted from cells using the GenElute™ Mammalian Total RNA Miniprep Kit (Sigma-Aldrich, RTN350). cDNA was reversely transcribed from 1 µg of RNA using the iScript cDNA synthesis kit (Bio-Rad, 1708891BUN) according to the manufacturer’s recommendations. Real-time quantitative PCR (RT-qPCR) was performed in three technical replicates per sample using SYBR Green master mix (Life Technologies, 11733046) on a ViiA 7 Real-Time PCR system (AB Applied Biosystems) utilizing specific primers at a concentration of 1 µM. Primer sequences used in this study are specified in **Supplementary Table 2**. Data analysis was performed with QuantStudio Real-Time PCR Software. CT values were detected for each sample and normalized to the housekeeping gene 60S Ribosomal Protein L19 (*RPL19*).

### RNA-seq library preparation

RNA sequencing (RNAseq) libraries were prepared using 1 µg of total RNA using the KAPA stranded mRNA-seq kit (Roche) according to the manufacturer’s specifications. KAPA-single index adapters (100 nM) (Roche, KK8702) were added to the A-tailed cDNA and the libraries underwent 10 cycles for amplification. Agentcourt AMPure XP beads (Beckman Coulter, A63880) were used for the 1x library clean-up. The fragment size of the library was assessed using the Agilent Bioanalyzer 2100 with the High Sensitivity DNA kit and was subsequently measured for the concentration by the High Sensitivity QuBit kit (Invitrogen, Q33230). Diluted libraries (4 nM) were pooled for sequencing on an Illumina HiSeq4000, aiming at 15-20 million SE50 reads per library.

### RNA-seq analysis

Trimming of adapters, polyA/T tails, and low-quality reads was performed using the Trimgalore! package (v0.6.4_dev) with default parameters. Using the Salmon package (v0.14.1) with default parameters^33^, the reads were aligned to the transcriptome and quantified using release 32 of the human reference transcriptome sequence and gene annotation. Counts were then imported into R using Tximport (v1.18.1). Differentially expressed genes (DEGs) were obtained using DEseq2 (v1.30.0) and were filtered to FDR-adjusted p-value < 0.05 and log2(Fold Change) > 1^34, 35^. Gene ontology enrichment was performed using Cluster Profiler (v3.18.1) and TTRUST was used to enrich transcription factors^36^. Shown heatmaps were generated using defined gene sets based on well described markers throughout literature and based on gene lists extracted from GO terms (**Supplementary Table 3 and Table 6**), displaying the z-score of TPM values across represented samples.

### Statistical analyses

In all the experiments, significant differences were determined by one-way analysis of variance (ANOVA), two-way ANOVA, adjusted for multiple comparisons, and two-tailed unpaired t-test. Data are presented as the mean ± standard error of the mean (SEM) or as fold change (FC). Data are visualized using GraphPad Prism 6 software (San Diego, CA, USA).

## RESULTS

### Sequential regulation of growth arrest and differentiation transcriptional programs upon hyperosmolar maintenance of liver models

To characterize the effects of hyperosmolarity on cell culture of hepatic models, we initially cultured the hepatoma cell line (HUH7.5), which resides in a proliferative, dedifferentiated, and immature state^37^, with standard culture media under plasma osmolarity (PlasOsm) levels (285-300 mOsm) and at increasing levels of hyperosmolarity. Two different organic osmolytes were employed to promote hyperosmolarity: an amino acid-based osmolyte, glycine (Osm^G^), previously shown to be present at high concentrations in the liver^29^, and a well-described inert sugar-based osmolyte, mannitol (Osm^M^), which has been widely used as the gold standard to establish HypOsm culture systems^38, 39^. Cell death was detected above 600 mOsm indicating that hepatoma cells reached toxic osmolarity levels (**Fig. 1A**). Following the measurement of the osmolarity within adult murine tissues, we show that the osmolarity of the liver ranges between 550-580 mOsm. Therefore, we considered that the *in vitro* osmolarity levels for hepatocytes using 250 mM glycine (Osm^G^) or mannitol (Osm^M^) supplementation (550 mOsm) resemble the physiological levels measured *in vivo* (**Fig. 1B**).At these HypOsm levels, no significant cell death was encountered (**Fig 1A**) which correlated with a cellular osmolar response defined by a significant nuclear accumulation of the osmolar master regulator NFAT5 in nearly 90% of the cells compared to cells cultured at PlasOsm levels (**Fig. 1C**). In addition, we detected an osmolar level- and time-dependent transcriptional upregulation of the osmo-protective NFAT5 target genes such as COX2 (*PTGS2*), Aldose Reductase (*AKR1B1*) and members of the Solute Carrier family (SLC) SMIT2 (*SLC5A3*) and BGT-1 (*SLC6A12*)^40^ (**Fig. S1A-B**). Neither the DNA damage marker pH2AX nor the cleaved Caspase-3 or apoptotic marker Annexin V were upregulated in cultured HUH7.5 cells in 550 mOsm Osm^G^ or Osm^M^ conditions, confirming the absence of HypOsm-induced cellular stress at this osmolar level (**Fig. 1D-F**). Therefore, we are able to categorize in vitro hyperosmolarity levels into healthy physiological and toxic ranges. Levels within physiological range (< 600 mOsm) induce a cellular osmolar response without affecting DNA damage in contrast to toxic osmolarity (> 600 mOsm) which induces cell death. Consequently, we considered 550 mOsm (Osm^G^, Osm^M^) to be within physiological HypOsm ranges as the optimal osmolar level to study the effects of prolonged HypOsm culture conditions on the cellular behavior of hepatic models.

**Figure 1:**
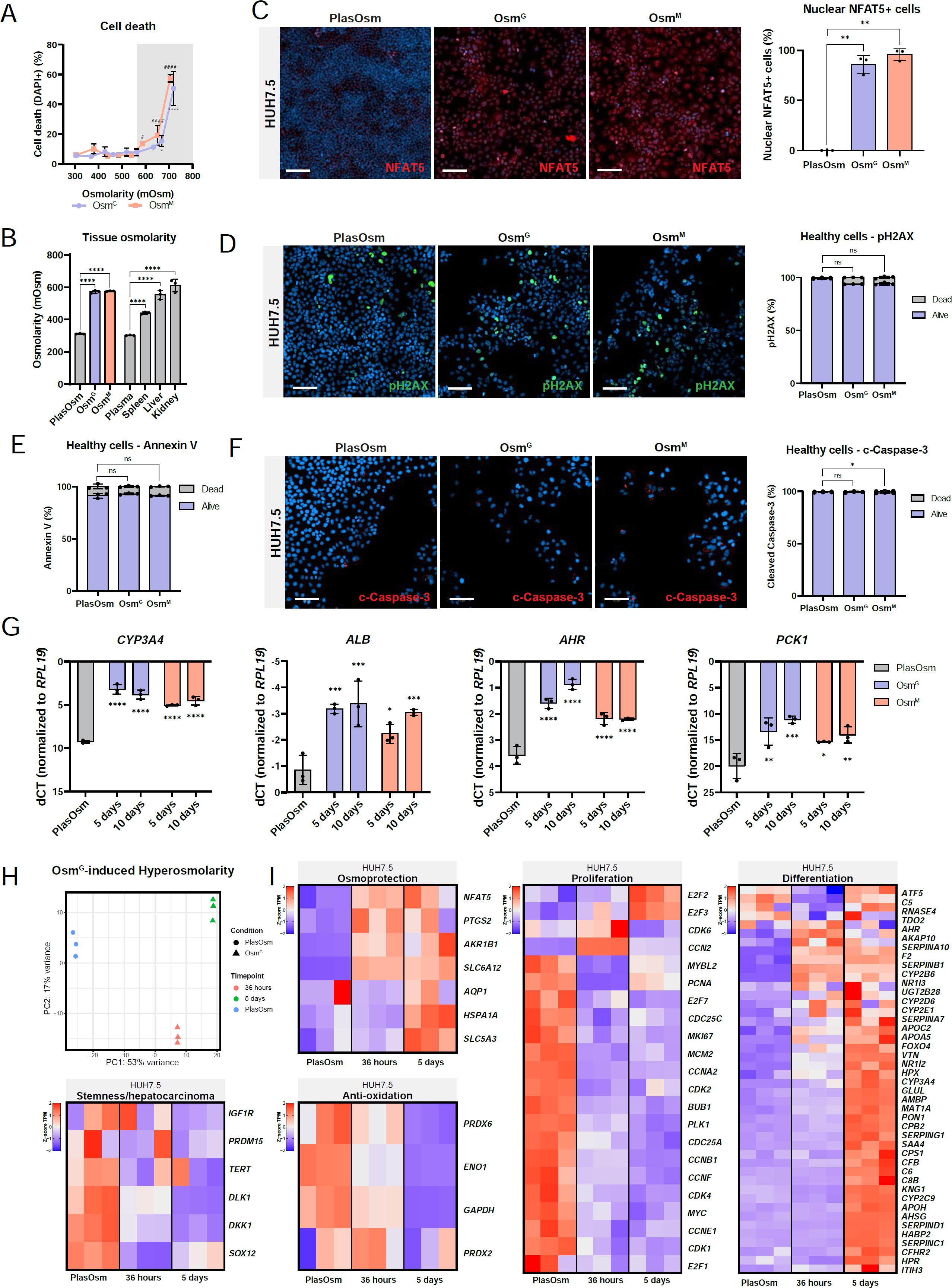
Sustained hyperosmolar exposure induces an adaptive osmolar response with enhanced cellular maturation.

**(A)** DAPI staining to quantify cell death with increasing Osm^G^ or Osm^M^ osmolarity values. Values with grey background are statistically significant compared to control. N=3, statistics performed by Brown-Forsythe and Welch ANOVA tests.
**(B)** Osmolarity of PlasOsm supplemented with either Osm^G^ or Osm^M^ next to osmolarity levels measured in adult murine plasma, spleen, liver, and kidneys and represented in mOsm. N=3, statistics by Ordinary one-way ANOVA
**(C)** Immunofluorescence staining (left) of NFAT5 upon Osm^G^ and Osm^M^ treatment in HUH7.5 cells with quantification of nuclear detection (right)N=3, statistics performed by Brown-Forsythe and Welch ANOVA tests. Scale Bar = 100μm.
**(D)** Immunofluorescence of DNA damage marker pH2AX in HUH7.5 cells upon 72h of Osm^G^ and Osm^M^ treatment. Scale bar = 100 µm. N=3, statistics by Kruskal-Wallis test.
**(E)** Annexin V staining of cells exposed to hyperosmolar stimuli after 72h of treatment. N=3, statistics performed by Brown-Forsythe and Welch ANOVA tests.
**(F)** Immunofluorescence of pre-apoptotic marker cleaved Caspase-3 after 72h of treatment of Osm^G^ and Osm^M^. Scale bar = 100 µm. N=3, statistics by Kruskal-Wallis test.
**(G)** Gene expression analysis of maturation genes *CYP3A4*, *ALB*, *AHR* and *PCK1* after 5 and 10 days of hyperosmolar treatment. Statistical significance was determined compared to the control condition. N=3, statistics by Two-way ANOVA.
**(H)** Principal Component Analysis of bulk RNA-seq analysis of PlasOsm and Osm^G^ treated HUH7.5 cells after 36 hours and 5 days of treatment.
**(I)** Heatmaps of bulk RNA-seq data representing the z-score of TPM values across presented sample conditions. Defined gene sets are summarized in Supplementary Table 3. All data represents mean ± SEM; *p < 0.05, **p < 0.01, ***p < 0.001, ****p <.0.0001.

Interestingly, when we prolonged HypOsm treatment of HUH7.5 cells for 5 and 10 days, a significant increase in the expression of mature hepatocyte markers (*CYP3A4*, *ALB*, *AHR,* and *PCK1* (**Fig. 1G**)) was observed in comparison to HUH7.5 cultured in PlasOsm conditions. This indicates that besides the functional adaptation to osmolar changes led by NFAT5 upregulation, HypOsm conditions might introduce a global transcriptional change resembling a more mature transcription profile. Transcriptome profiling by RNA-seq was used to identify the landscape of early and later events 36 hours and 5 days following physiological HypOsm (Osm^G^) treatment (**Fig. 1H**). To evaluate the transcriptional changes, we defined clear gene sets that consist of widely used markers throughout literature to represent the cellular processes emerging from our analysis (**Supplementary Table 3**). Using the stringent criteria (FDR < 0.05, log2(fold change) > 1), 1830 and 2471 differentially expressed genes were upregulated and 1177 and 1852 were downregulated after 36h and 5 days of treatment, respectively (**Supplementary Table 4**). Among the upregulated genes, we identified osmo-protective genes including members of the solute carrier family (*SLC5A3*, *SLC6A12*), while among the downregulated genes, we identified genes involved in mitochondrial oxidation such as *PRDX6*, *ENO1,* and *GAPDH* in line with the notion that HUH7.5 cells adaptively responded to physiological hyperosmolarity (**Fig. 1I**). Interestingly, Gene Ontology (GO) term enrichment analysis at 36 hours revealed that among the downregulated genes, cell cycle-related terms were highly significant enriched (**Fig. S1C, Supplementary Table 5**), indicated by the downregulation of the pro-proliferative (*CDK1*, *CDK2, E2F1, MYC*) and proliferative markers (*MKI67*, *PCNA*). This was maintained following 5 days of Osm^G^ culture (**Fig. 1I**). GO analysis identified metabolic terms as highly significantly enriched at day 5 (**Fig. S1D, Supplementary Table 5**), including genes involved in drug metabolization such as *NR1I2/3* and CYP450 enzymes, all known to be expressed in mature hepatocytes cells^41^. Interestingly, most of the maturation genes were only upregulated after 5 days but not after 36 hours of Osm^G^ culture, indicating that inhibition of the cell cycle precedes induction of maturation when HUH7.5 cells are cultured under physiological HypOsm conditions.

These findings showed that in vitro physiological HypOsm culture conditions induced a hyperosmotic adaptive response without triggering cell stress or apoptosis. Additionally, we demonstrated that prolonged HypOsm treatment induced a transcriptional rewiring that goes further than specific HypOsm gene transcriptional response, characterized by a sequential downregulation of cell cycle genes and induction of maturation genes.

### Hyperosmolarity induces a G_0_ cell cycle arrest in a p53-independent manner

As transcriptional alterations in the expression of cell cycle regulators were observed at early time points following physiological HypOsm treatment (**Fig. 1**), we tested the effect of HypOsm culture on hepatoma cell expansion/proliferation. Cell counting and EdU uptake experiments showed that HUH7.5 cells cultured in Osm^G^ or Osm^M^ conditions caused a significant cell cycle arrest already after 24 hours compared to cells cultured in PlasOsm conditions (**Fig. 2A, 2B**). To assess cell cycle dynamics, we introduced the fluorescent FUCCI reporter construct into the HUH7.5 cells^42^ (**Fig. S2A**). Cells cycling through the G_0_/G_1_ phase are observed in red by CDT1 excitation, and S/G_2_ cells express fluorescent protein geminin^43^. While in PlasOsm culture medium cells displayed a default FUCCI profile highly enriched in cells in S/G_2_ (60%), Osm^G^ or Osm^M^ treatment resulted in a significant enrichment of cells in the G_0_/G_1_ cell cycle phase (**Fig. 2C**). Cells express KI67 during proliferating cell cycle phases but not during the G_0_ phase, indicating that cells that lose KI67 expression undergo G_0_ cell cycle exit^44^. Under PlasOsm conditions, 90% of the HUH7.5 cells resided in a proliferative (KI67^+^) state. In contrast, Osm^G^-treated cells became predominantly KI67 negative demonstrating growth arrest and accumulation of cells in the G_0_ phase (**Fig. 2D**). This was further confirmed by the visualization of KI67 within the G_0_/G_1_ cells of the FUCCI reporter line (**Fig. 2E**). Cell cycle arrest at G_0_/G_1_ was considerably present at concentrations higher than 450 mOsm reaching the highest percentage of arrested cells at concentrations around 600 mOsm. Increasing osmolar levels at toxic osmolarity ranges above 600 mOsm reduced the number of cells in G_0_/G_1_, potentially explained by the concomitantly significant increase in cell death visualized by DAPI staining (**Fig. 2F**).

**Figure 2:**
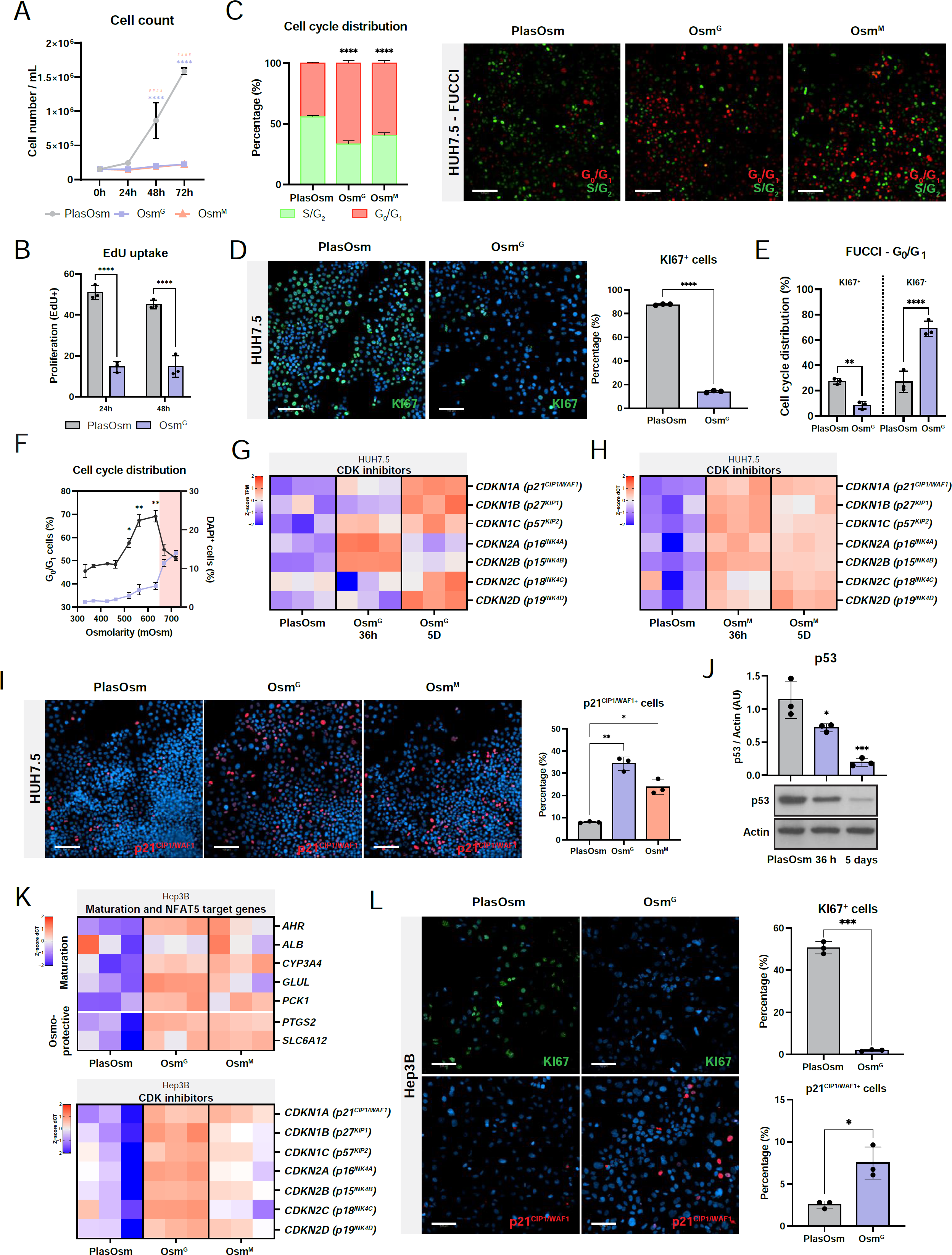
Hyperosmolarity induces a G_0_-mediated cell cycle arrest in a p53-independent manner.

**(A)** Cell counts of HUH7.5 cells in plasma osmolar and HypOsm conditions after 24h, 48h and 72h. N=6, statistics by Two-way ANOVA with Bonferroni correction.
**(B)** EdU uptake of HUH7.5 cells after 24h and 48h of PlasOsm or HypOsm culture conditions. N=3, statistics by Two-way ANOVA with Bonferroni correction.
**(C)** HUH7.5 cells containing the FUCCI reporter after 72 hours of HypOsm conditions, visualized by fluorescent microscopy (right) and quantified to show cell cycle distribution (left). Scale bar = 100 µm. N=3, statistics by Two-way ANOVA.
**(D)** Representative immunofluorescent staining (left) and quantification (right) of KI67 after 24 hours of OsmG treatment. N=3, statistics by Student’s t-test.
**(E)** Using the FUCCI reporter, cells were quantified and co-stained with KI67. N=3, statistics by Two-way ANOVA with Bonferroni correction.
**(F)** Cell cycle distribution at different osmolarities using the FUCCI reporter and correlation with cell death stained by DAPI in live cells. N=3. DAPI data under Osm^G^ from Figure 1A. Statistics for G_0_/G_1_ cells were performed by Brown-Forsythe and Welch ANOVA tests, all osmolarity values compared to control.
**(G)** Gene expression analysis of CDK inhibitors after 36 hours and 5 days of treatment with Osm^G^ visualized in z-score of the TPM values. N=3.
**(H)** Gene expression analysis of CDK inhibitors after 36 hours and 5 days of treatment with Osm^M^ normalized to *RPL19* expression and visualized in z-score of the dCT values. N=3.
**(I)** Representative fluorescent staining (left) and quantification (right) of p21^CIP1/WAF1^ after 24 hours of Osm^G^ and Osm^M^ treatment. Scale bar = 100 µm. N=3, statistics by Brown-Forsythe and Welch ANOVA tests.
**(J)** Representative western blot (down) and quantification (up) of p53 and Actin after 36 hours and 5 days of Osm^G^ treatment. N=3, statistics by Brown-Forsythe and Welch ANOVA tests.
**(K)** Gene expression analysis of hepatic maturation markers (top), NFAT5 target genes (middle) and CDK inhibitors (bottom) in Hep3B cells upon Osm^G^ and Osm^M^ treatment after 5 days of hyperosmolar treatment. Gene expression is normalized to housekeeping gene RPL19. Data is visualized in z-score of the dCT levels N=3.
**(L)** Immunofluorescent staining of KI67 and p21CIP1/WAF1 in Hep3B cells treated with Osm^G^ for 24 hours. Scale bar = 100 µm. N=3, statistics by Welch’s t-test. All data represents mean ± SEM; *p < 0.05, **p < 0.01, ***p < 0.001, ****p <.0.0001.

As previous reports have shown that the liver is enriched in several amino acids (AAs), including glycine, alanine, or serine, we investigated whether other AAs with osmolyte activity might mirror the effect of glycine on the cell cycle^45, 46^. Indeed, as several distinct AAs induced a cell cycle arrest without triggering cell death, this implies that the cell cycle exit is a shared cellular response to the HypOsm treatment across several organic osmolytes (**Fig. S2B**).

Given the importance of CDKIs in growth arrest, we assessed the expression levels of CDKIs following exposure to Osm^G^ or Osm^M^. The INK4 family members p15^INK4B^ (*CDKN2B*), p18^INK4C^ (*CDKN2C*), p19^INK4D^ (*CDKN2D*), and the members of the CIP/KIP family p21^CIP1/WAF1^ (*CDKN1A*) and p27^KIP1^ (*CDKN1B*) and p57^KIP2^ (*CDKN1C*) were significantly upregulated upon Osm^G^ or Osm^M^ exposure of HUH7.5 cells for 36 hours and 5 days (**Fig. 2G-H**). Immunofluorescence analysis confirmed a significant upregulation of p21^CIP1/WAF1^, one of the main proteins involved and responsible for cell cycle exit ^47^ (**Fig. 2I**).

In mammals, p53 is the master regulator of cell cycle arrest by tightly controlling the expression of *CDKN1A*. Additionally, p53 is increased in toxic osmotic culture conditions and regulates cell differentiation by association with stem cell differentiation genes^48–53^. We detected increased p53 expression under toxic HypOsm (650 mOsm) levels in agreement with previous reports^48, 50, 51^ (**Fig. S2C**). Surprisingly, p53 was significantly decreased at protein level upon physiological Osm^G^ treatment, suggesting that the physiological HypOsm conditions induced the upregulation of CDKi expression, resulting in cell cycle exit in a p53-independent manner (**Fig. 2J**). To confirm whether p53 is dispensable for the HypOsm-induced cell cycle arrest, we treated the p53-null hepatocarcinoma cell line Hep3B with physiological HypOsm culture conditions. Osm^G^ or Osm^M^ treatment of Hep3B induced the expression of osmo-protective genes, CDKi *CDKN1A,* and maturation markers while downregulating proliferation marker KI67, in line with the results obtained with the HUH7.5 cell line (**Fig. 2K** and **2L**). These findings suggest that the mechanisms to facilitate the cell cycle exit in response to physiological HypOsm conditions take place in a p53-independent manner.

### iPSC-derived hepatocyte-like cells respond to increased osmolarity inducing growth arrest and cell maturation

Although PSC differentiation to cells with hepatocyte features has been well described, most protocols generate cells resembling a fetal and immature differentiation state^54–58^. Indeed, a transcriptional meta-analysis of publicly available data^29^ comparing hiPSC-derived hepatocyte-like cells (hiPSC-HLCs) to cultured mature primary human hepatocytes (PHH) revealed that hiPSC-based hepatic progeny remained in an immature state as indicated by the low expression of metabolic GO processes and displayed a proliferative transcriptional profile (**Fig. S3A, Supplementary Table 6**).

Inspired by our findings in the hepatoma line, we subsequently aimed to understand whether physiological HypOsm conditions could improve the maturation and growth arrest of hiPSC-HLC. To obtain hiPSC-derived HLCs, we used a previously described protocol employing liver differentiation medium (LDM) in PlasOsm conditions supplemented with a cascade of growth factors^31, 59^ (**Fig. S3B**). When undifferentiated hiPSCs were exposed to HypOsm conditions, these instantly underwent apoptosis within 24 hours (**Fig. 3A**), indicating that pluripotent hiPSCs cannot adapt to elevated HypOsm challenges (550 mOsm). This is in line with previous reports where cultured embryos were sensitive to low HypOsm levels^60^. In contrast, RNA-seq analysis of hepatoblast-like cells obtained on day 12 of the HLC differentiation protocol revealed a significant upregulation of SLC family genes compared to hiPSC cells, suggesting that lineage-committed cells, unlike hiPSC, are equipped with the proper transport systems to adapt to a higher osmolar challenge (**Fig. S3C**). Therefore, we started treatment of hiPSC progeny from the hepatoblast stage of the differentiation protocol (Day 12, **Fig. S3B**) with Osm^G^ or Osm^M^, as these did not induce the cleavage of apoptotic marker caspase-3 (**Fig. 3B**).

**Figure 3:**
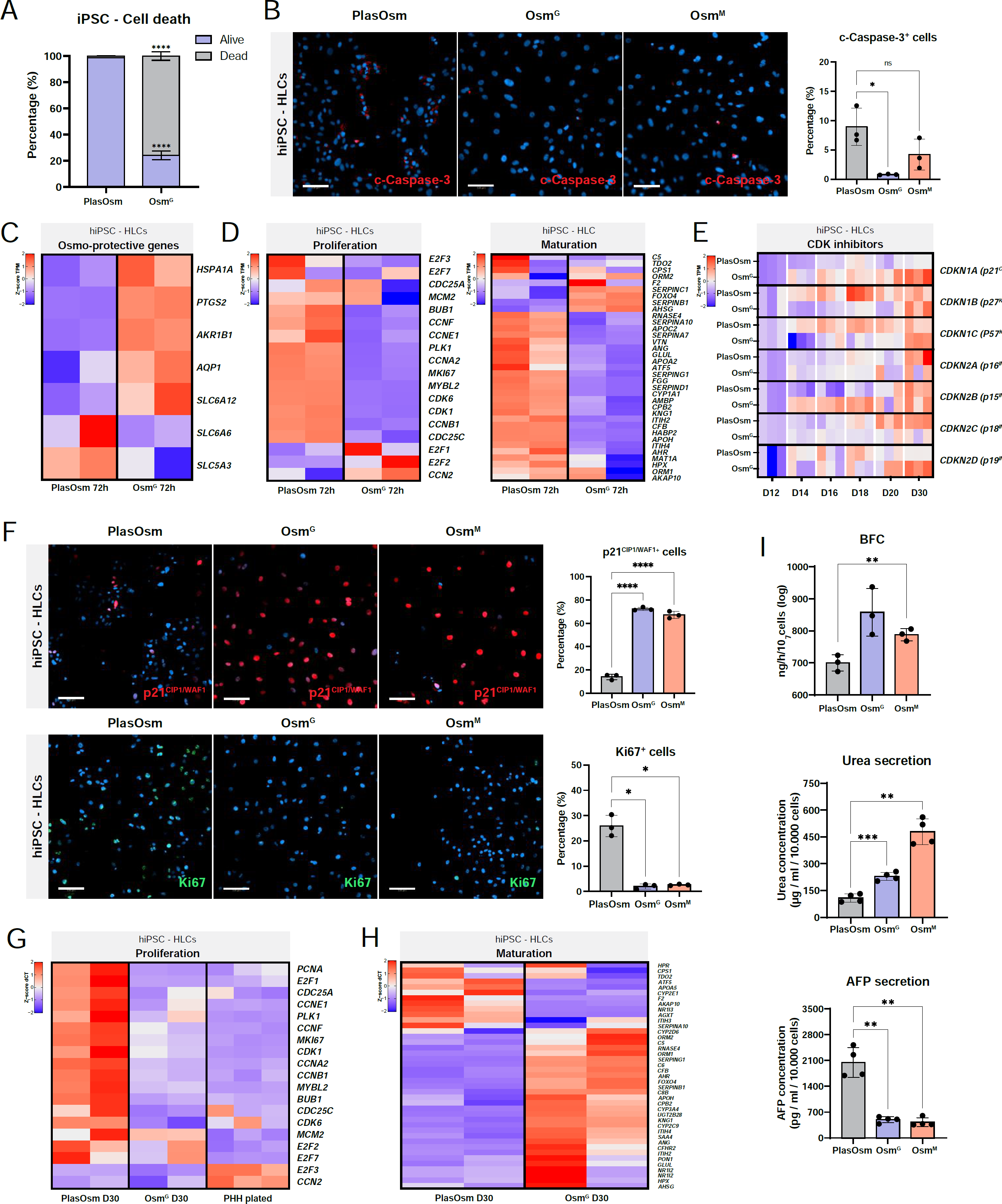
Hyperosmolarity-induced growth arrest and lineage maturation in hiPSC-derived HLCs.

**(A)** Viability by DAPI staining of hiPSC cultured in pluripotent conditions 24 hours after Osm^G^ treatment. N=3. Statistics by Two-way ANOVA with Bonferroni correction.
**(B)** Representative image of fluorescent detection (left) and quantification (right) of cleaved-caspase 3 after 72h of Osm^G^ treatment. N=3. Statistics by Kruskal-Wallis test.
**(C)** Heatmap of genes involved in osmo-protection of hiPSC derived HLCs 72h after Osm^G^ treatment visualized in z-score of the TPM values.
**(D)** Heatmap of genes involved in proliferation (left) and hepatic maturation (right) 72h after Osm^G^ treatment visualized in z-score of the TPM values.
**(E)** Heatmap of CDK inhibitors analyzed by RT-PCR at different timepoints after Osm^G^ treatment represented in z-score of dCT values.
**(F)** Representative images of fluorescent staining (down) and quantification (up) of KI67 and p21^CIP1/WAF1^ after 72h of Osm^G^ treatment. Scale bar = 100 µm. N=3, statistics by Brown-Forsythe and Welch ANOVA tests.
**(G)** Heatmap of proliferation genes after Osm^G^ treatment in hiPSC derived HLCs at D30 of differentiation compared to plated PHHs.
**(H)** Heatmap of hepatic maturation markers upon Osm^G^ treatment in hiPSC derived HLCs at D30 of differentiation.
**(I)** Functional comparison between HLC control cells, cells treated with Osm^G^ and Osm^M^ medium for BFC metabolism, urea secretion and AFP secretion. Statistics by Brown-Forsythe and Welch ANOVA tests. All data represents mean ± SEM; *p < 0.05, **p < 0.01, ***p < 0.001, ****p <.0.0001.

Following treatment with Osm^G^, the transcriptional changes were assessed by RNA-seq after 72 hours of exposure and on day 30 of the iPSC-HLC differentiation protocol to capture early and late cellular changes. Using stringent criteria (FDR < 0.05, log2(fold change) > 1), we identified 1800 up- and 2832 downregulated DEGs after 72 hours of treatment and 1898 up- and 3073 downregulated DEGs were identified on day 30 of the differentiation protocol (**Supplementary Table 7**). Osmo-protective genes such as *AQP1* and *SLC6A12* were upregulated after 72h as a direct response to the hyperosmolar treatment (**Fig. 3C**). At day 30 of the differentiation under HypOsm conditions, gene ontology enrichment of DEGs between Osm^G^ and PlasOsm differentiation identified enrichment of cell proliferative terms in the downregulated DEGs, while cell metabolic processes were enriched in the upregulated DEGs. (**Fig. S3D, Supplementary Table 8**). In agreement with our previous results in hepatoma cell lines, HypOsm culture conditions during hiPSC-HLC differentiation revealed downregulated proliferation markers early upon treatment, while maturation markers were not yet upregulated upon 72h of HypOsm conditions, indicating that cells focus their early transcriptional response to HypOsm conditions on regulating their cell cycle exit (**Fig. 3D**).

Under HypOsm conditions, RT-qPCR analysis demonstrated an increase in the elevated levels of the CDKis *CDKN1A* and *CDKN2B* already after 72h, increasing further throughout days 20 and 30 (**Fig. 3E**). Furthermore, we observed a significant upregulation of the percentage of p21^CIP1/WAF1^ positive cells together with a compelling decrease of the proliferation marker KI67 in Osm^G^ and Osm^M^ culture conditions compared to PlasOsm conditions (**Fig. 3F**). Co-detection of p21^CIP1/WAF1^ and p53 in the cells after 72 hours of hyperosmolar treatment showed that the majority of p21^CIP1/WAF1^ positive cells were lacking p53 expression, confirming our findings for the HUH7.5 cells and suggesting that p21 expression and cell cycle exit appear to be p53 independent (**Fig. S3E**). Interestingly, unlike PlasOsm cultured HLCs, the Osm^G^-induced HLC cell proliferation signature was similar to that of plated PHH obtained from publicly available data (**Fig. 3G**). As we had seen for HUH7.5 cells, prolonged maintenance of physiological hyperosmolar conditions throughout the differentiation protocol (day 12 to day 30) caused a transcriptional increase in mature hepatic cell marker genes suggesting that HypOsm-induced maturation may be a consequence of the preceding growth arrest early during HypOsm treatment (**Fig. 3H, S3F**). This transcriptional profile was also translated in the functional improvement of the HLCs, demonstrated by significantly increased CYP3A4-dependent BFC metabolism, urea secretion, and the reduced fetal AFP secretion in HypOsm (Osm^G^ and Osm^M^) versus PlasOsm treated HLCs (**Fig. 3I**).

Altogether, our results demonstrate that physiological HypOsm culture conditions induced a growth arrest and improved maturation of differentiating hiPSC-HLCs. Furthermore, the transcriptional cell cycle signature of Osm^G^ and Osm^M^-treated hiPSC-HLCs resembled that of cultured PHH. Although a number of mature hepatocyte marker genes were significantly higher expressed in Osm^G^ conditions compared with PlasOsm HLCs, which was reflected in functional improvement, expression of other mature marker genes in OsmG-treated hiPSC-HLCs remained lower than in plated PHH, indicating that several factors are still lacking to further drive hPSC-HLCs towards their *in vivo* counterpart (**Fig. S3G**).

### Hyperosmolarity-induced WNT inhibition regulates transcriptional cell maturation but not growth arrest

To gain insight into the signaling pathways underlying the HypOsm-induced growth arrest and transcriptional maturation, we evaluated the RNA-seq analysis using more stringent parameters (FDR < 0.05, log2(fold change) > 1.5). Among the differentially regulated signaling pathways, WNT signaling appeared to be significantly differentially regulated (**Figure S4A, Supplementary Table 9**). This signaling pathway has been described as an indispensable pathway in development and regeneration, regulating the cell cycle, stemness, and differentiation^61^. In line, the protein levels of non-phosphorylated (active) β-catenin, the key co-factor regulating canonical WNT transcriptional gene expression^61^, were markedly downregulated after Osm^G^ treatment, indicating that Osm^G^ induces a transcriptional suppressed state of WNT signaling (**Figure S4B**).

To investigate whether HypOsm-induced WNT suppression is necessary for the induction of cell growth arrest or cell maturation, we activated WNT signaling by treating HUH7.5 cells with the specific GSK3 inhibitor CHIR99021 (WNT activator; CHIR). CHIR treatment induced transcriptional WNT activation under HypOsm conditions as indicated by the WNT responsive 7TGP reporter and expression levels of its target genes^62^ (**Fig. S4C-D**). Principal component analysis of RNA-seq samples revealed that CHIR-treated samples shifted downwards along the PC2 axis, which accounted for genes involved in cell differentiation (**Fig. S4E, Supplementary Table 10**). In agreement, CHIR treatment reduced the expression of some specific maturation and differentiation genes (**Fig. S4F**). Interestingly, key hepatoma genes such as *SOX12* were upregulated when WNT was reactivated under Osm^G^ conditions, suggesting that WNT might control the bridge between stemness and maturation (**Fig. S4F**).

Although the WNT pathway has been described to be involved in cell proliferation^63–65^, WNT activation by CHIR in HypOsm conditions did not affect proliferative markers at the RNA level and was unable to rescue the HypOsm-induced cell cycle arrest visualized by KI67 staining in HUH7.5 cells (**Fig. S4G-I**). Hence, WNT activation did not restore the cell cycle exit, but reverted the transcriptional maturation profile under hyperosmolar conditions.

We next assessed whether the WNT pathway was also involved in the HypOsm-induced hiPSC-based HLC maturation. Of note, the transcriptional meta-analysis of hiPSC-derived HLCs compared with plated PHHs revealed significantly increased WNT signaling in HLCs, suggesting that WNT suppression might induce HLC maturation (**Fig. S4J**). As in the HUH7.5 model, hyperosmolar conditions during hiPSC differentiation induced reduced levels of active β-catenin concomitantly with increasing levels of the maturation marker CYP3A4 (**Fig. 4A**). When HypOsm cultures were also treated with CHIR to reactivate WNT signaling, gene expression of CYP3A4 together with other maturation markers was significantly decreased in reverse correlation with activation of Wnt target genes (**Fig. 4B, S4K**). We further verified the suppressive effects of WNT on maturation as seen with CYP3A4 expression at the protein level (**Fig. 4C**). As shown in HUH7.5 cells, WNT reactivation had minimal effects on the expression levels of genes involved in the HypOsm-induced cell cycle arrest such as *CDKN1A* and KI67 at the protein level in hiPSC-derived HLCs differentiation (**Fig. 4D-E, S4L**).

**Figure 4:**
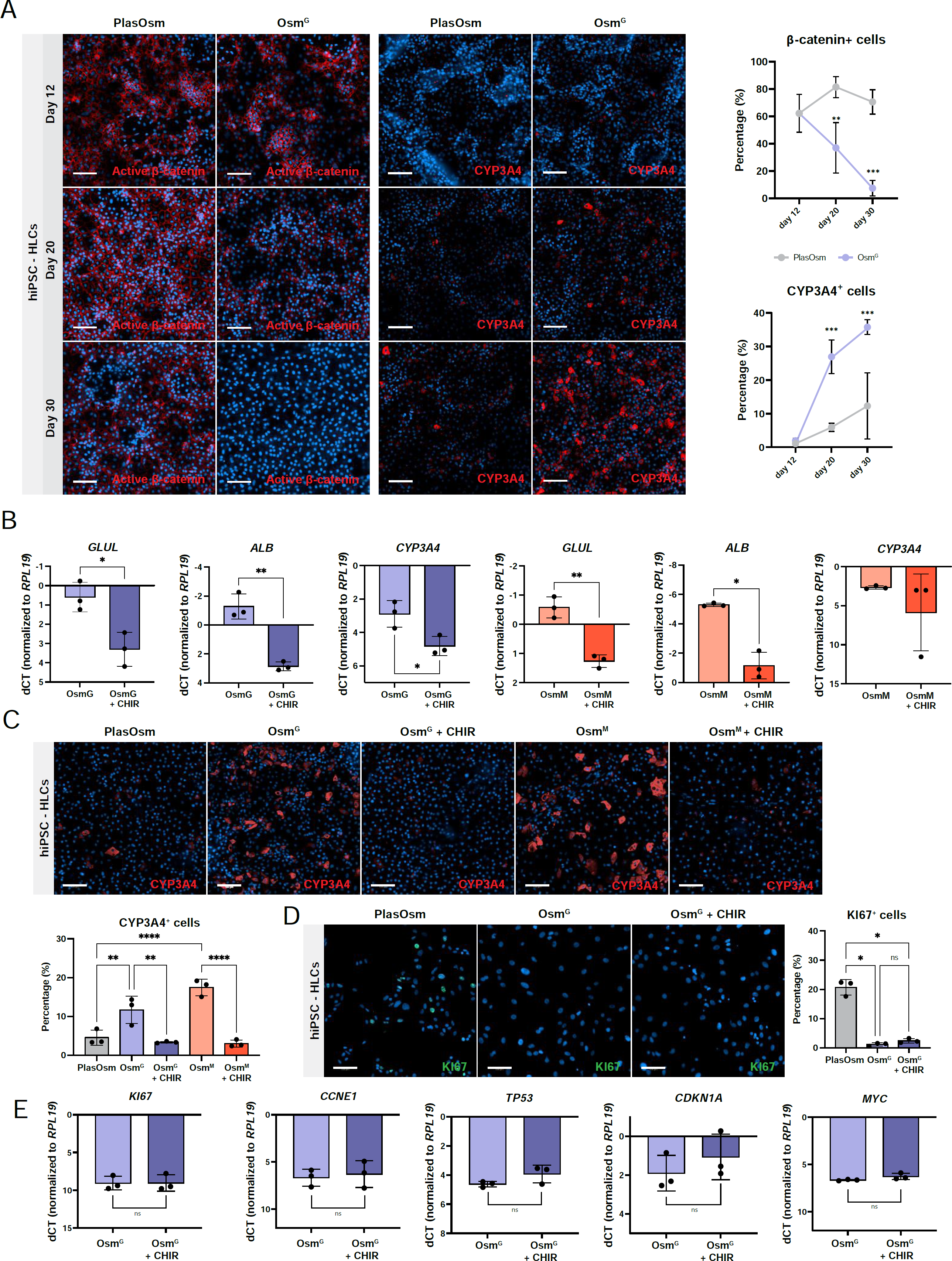
Wnt signaling regulates lineage specification during hyperosmolarity guided differentiation.

**(A)** Immunofluorescence (left) and quantification (right) of active β-catenin and CYP3A4 at day 12, 20 and 30 of the hiPSC-HLC differentiation protocol. Scale bar = 100 µm. N=3, statistics by Two-way ANOVA.
**(B)** Day 30 gene expression of *GLUL*, *ALB* and *CYP3A4* genes of hiPSC-HLC differentiation samples treated with Osm^G^ (in blue) or Osm^M^ (in red) with and without CHIR99021 from day 12 to day 30. N=3, statistics by Welch’s t-test.
**(C)** Representative immunofluorescence images (up) and quantification (down) of CYP3A4 at D30 of hiPSC derived HLC differentiation protocol, cells treated with Osm^G^ or Osm^M^ with and without CHIR99021. Scale bar = 100 µm. N=3, statistics by Two-way ANOVA.
**(D)** Representative immunofluorescence images (left) and quantification (right) of KI67 at D30 of hiPSC-derived HLC differentiation protocol, cells treated with Osm^G^ with and without CHIR99021. Scale bar = 100 µm. N=3, statistics by Two-way ANOVA.
**(E)** Day 30 gene expression of proliferation genes of hiPSC-derived HLC samples treated with Osm^G^ with and without CHIR99021. N=3, statistics by Welch’s t-test. All data represents mean ± SEM; *p < 0.05, **p < 0.01, ***p< 0.001, ****p<0.0001.

Our findings suggest that transcriptional WNT suppression takes place downstream of the HypOsm conditions, indicating that transcriptional WNT repression is favorable for hepatocyte differentiation. In agreement, the rescue of WNT repression by pharmacological GSK3 inhibition counteracted the increased expression of hepatocyte maturation genes. However, WNT reactivation did not affect cell cycle gene expression in HypOsm-treated hepatoma and differentiating PSC cells. This demonstrates that HypOsm-mediated WNT inhibition is responsible for the HypOsm-mediated transcriptional maturation, but not cell cycle arrest.

### NF-κВ orchestrates cellular maturation via a hyperosmolarity-induced cell cycle exit

To assess how hyperosmolarity affects cell cycle changes, we searched for other pathways that were modified in HypOsm-dependent conditions. Using the regulatory network-based transcription factor prediction tool TTRUST on identified DEGs, the NF-κВ pathway appeared to be differentially regulated upon HypOsm exposure of HUH7.5 cells. Interestingly, the transcription factor prediction of DEGs at 36 hours captured members of the Rel family including NF-κВ members as possible regulators during the early events taking place upon hyperosmolar treatment (**Figure S5A, Supplementary Table 11**). Previous studies reported developmental defects in mouse and impaired differentiation of human PSCs upon the deletion of NF-κВ components^60, 66, 67^, highlighting the essential role of the NF-κВ signaling pathway in pluripotent differentiation. In line, the RNA-seq analysis identified low levels of NF-κВ target genes in hiPSC-HLCs cultured under PlasOsm compared with plated PHHs (**Fig. S5B**).

To assess whether NF-κВ signaling becomes activated in hiPSC-HLCs and hepatoma line HUH7.5 under HypOsm conditions, cells were stained for the inhibitor of NF-κВ signaling κВα (IκВα) which retains NF-κВ subunits in the cytoplasm, rendering the pathway inactive. hiPSC-HLCs and hepatoma cells cultured in PlasOsm conditions showed high levels of IκВα expression, indicating that PlasOsm culture conditions maintain low NF-κВ activity levels (**Fig. 5A-B** and **S5C**). In contrast, HypOsm conditions significantly decreased IκВα expression with a concomitant fast induction of the direct NF-κВ target, ICAM-1 (**Fig. 5A-B** and **S5C**). Moreover, the expression of NF-κВ target genes was significantly increased in HypOsm-treated hiPSC-HLCs, indicating that these conditions activate NF-κВ (**Fig. S5D**).

**Figure 5:**
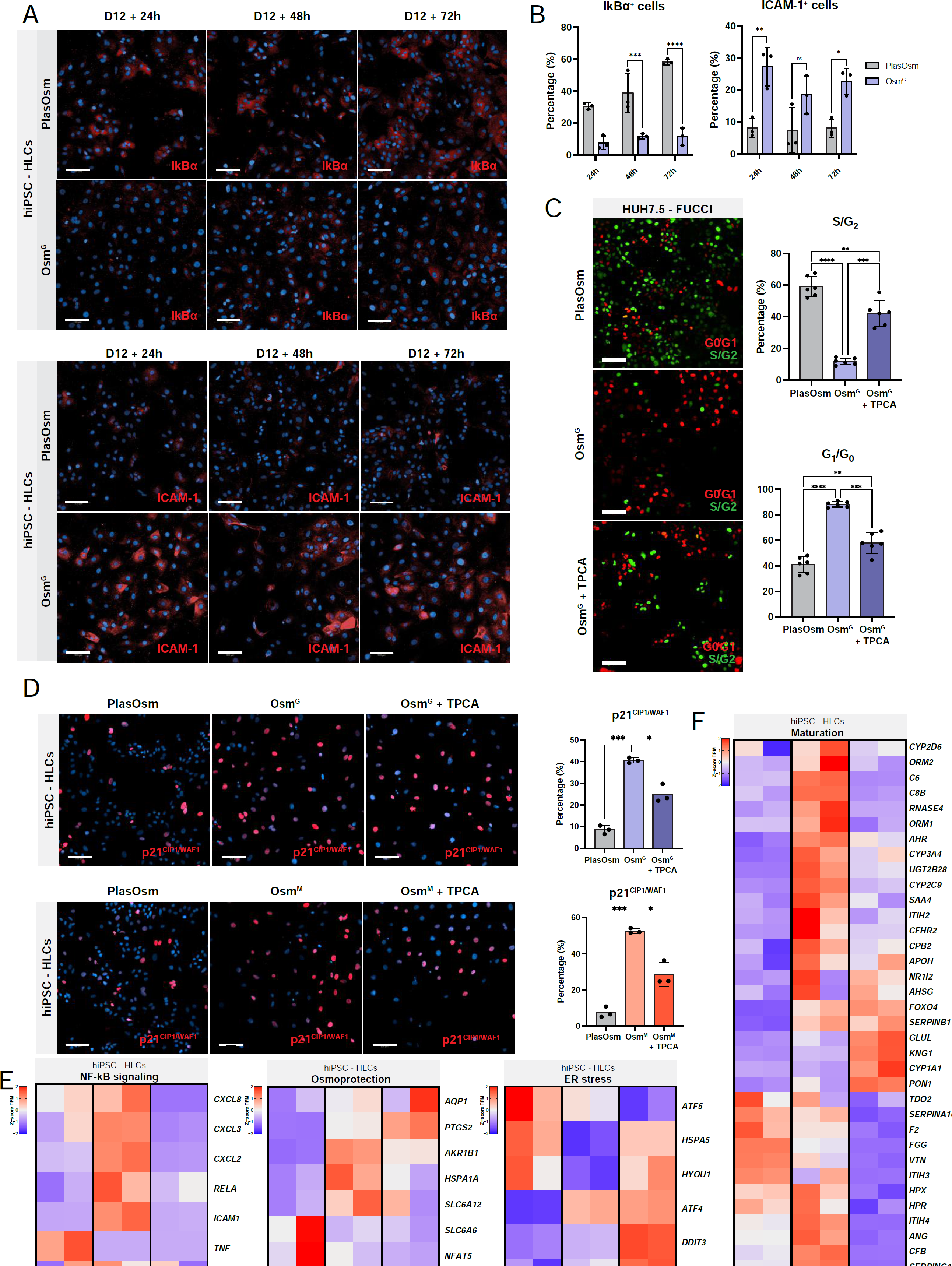
Hyperosmolarity leads to NF-κВ-mediated cell cycle arrest.

**(A)** Representative immunofluorescence images of IκВα and ICAM-1 at D12+24 hours, D12+48 hours and D12+72 hours of Osm^G^ treatment in hiPSC-derived HLCs. Scale bar = 100 µm.
**(B)** Quantification of immunofluorescent detection of IκВα and ICAM-1 from Figure (A). N=3, statistics by Two-way ANOVA.
**(C)** Representative fluorescent images (left) and quantification (right) of FUCCI-HUH7.5 cells under PlasOsm, Osm^G^ and Osm^G^ with TPCA-1 culture conditions for 72 hours. Scale bar = 100 µm. N=3, statistics by Brown-Forsythe and Welch ANOVA tests.
**(D)** Representative immunofluorescence images (left) and quantification (right) of p21^CIP1/WAF1^ under PlasOsm, HypOsm or TPCA-1 medium, as indicated, on day 12 + 72h of the hiPSC-derived HLCs protocol. Scale bar = 100 µm. N=3, statistics by Brown-Forsythe and Welch ANOVA tests.
**(E)** Heatmaps of bulk RNA-seq data representing the z-score of TPM values across presented sample conditions in hiPSC-HLCs showing genes associated with NF-κВ signaling, osmo-protection and ER stress.
**(F)** Heatmap of bulk RNA-seq data representing the z-score of TPM values across presented sample conditions in hiPSC-HLCs depicting genes associated with hepatic differentiation. All data represents mean ± SEM; *p < 0.05, **p < 0.01, ***p< 0.001, ****p<0.0001.

We next treated HypOsm-cultured cells with the IKK2 inhibitor TPCA-1 to suppress NF-κВ activity^68^. Using the FUCCI reporter system in HUH7.5 cells, we found that TPCA-1-mediated NF-κВ inhibition ablated the HypOsm-induced cell cycle inhibition, where cells now again accumulated in the S/G_2_ phase and significantly reduced the proportion of cells in G_0_/G_1_, mimicking the pattern seen for HUH7.5 cells cultured in PlasOsm conditions (**Fig. 5C**). Additionally, we also found a reversion of the induction of the growth arrest marker p21^CIP1/WAF1^, upon TPCA-1 treatment in both hiPSC-HLCs and HUH7.5 cells (**Fig. 5D** and **S5E**). Transcriptional analysis of RNA-seq data of hiPSC-HLCs cultured in Osm^G^ with TPCA-1 revealed that osmo-protective genes such as *AKR1B1*, *HSPA1A*, *SLC6A12,* and *SLC5A3* were reduced upon NF-κВ inhibition together with several NF-κВ target genes (**Fig. 5E, S5F-G, Supplementary Table 12 and Table 13**). Furthermore, GO enrichment revealed a remarkable collection of responses to misfolded proteins and ER stress, attributing to cellular stress and apoptosis upon TPCA-1-induced NF-κВ inhibition under HypOsm conditions (**Fig. 5E** and **S5H-I, Supplementary Table 14**). Consequently, the remaining cells failed to upregulate maturation markers and lost their differentiation signature (**Fig. 5F**).

Collectively, our findings indicate that NF-κВ controls hyperosmotic protection and growth arrest, affecting maturation and differentiation quality under hyperosmolarity. Furthermore, NF-κВ signaling has proven to be essential for the cellular survival and differentiation integrity, as inhibition of NF-κВ during HypOsm treatment affects the viability of the cells due to ER stress, compromising their differentiation.

### Prolonged hyperosmolarity is necessary for the stabilization of growth arrest and transcriptional maturation

The maintenance of both hepatic models in HypOsm culture conditions induced the improvement of both maturation and growth arrest. Once cells exit the cell cycle, they may either reside in a permanent active arrest, known as senescence; or in a reversible growth arrest, known as quiescence, from which cells resume proliferation. Senescence generally occurs in the G_1_ and in some cases the G_2_ phase of the cell cycle as opposed to quiescence taking place in the G_0_ phase, suggesting that HypOsm-arrested hepatic cell models might reside in a quiescent state^69^.

When HUH7.5 cultures treated with HypOsm culture conditions were transferred back to the PlasOsm range (Rev^G^ and Rev^M^; **Fig. 6A**), the levels of the osmo-protective genes *AKR1B1* and *PTGS2* were reduced indicating that cells remained sensitive to reversibly modulating the osmolar conditions. Concomitantly, levels of the CDKi p21^CIP1/WAF1^ (*CDKN1A*) and p57^KIP2^ (*CDKN1C*) were also reduced at the transcriptional level, suggesting that cells re-entered the cell cycle and lost their adaptive osmolar response (**Fig. 6B**). Moreover, together with the loss of growth arrest, the previous enhancement of maturation in the hepatoma line was significantly reduced as shown by decreased expression of maturation markers *CYP3A4*, *ALB*, *GLUL* and *PCK1* (**Fig. 6C**). We confirmed a reduced expression of the maturation marker CYP3A4 and the cell cycle inhibitor p21^CIP1/WAF1^ at the protein level (**Fig. 6D**). This cell cycle re-entry was also seen by an increased cell number at the end of the treatment (**Fig. 6E**). Interestingly, underlying these phenotypical changes, the previously dissected NF-κВ and WNT signaling pathways were affected accordingly as the osmolar reversion led to the inactivation of the NF-κВ pathway, indicated by the reduction in ICAM-1, while WNT signaling activity was restored as shown by the presence of β-catenin in its non-phosphorylated (active) form (**Fig. 6D**).

**Figure 6:**
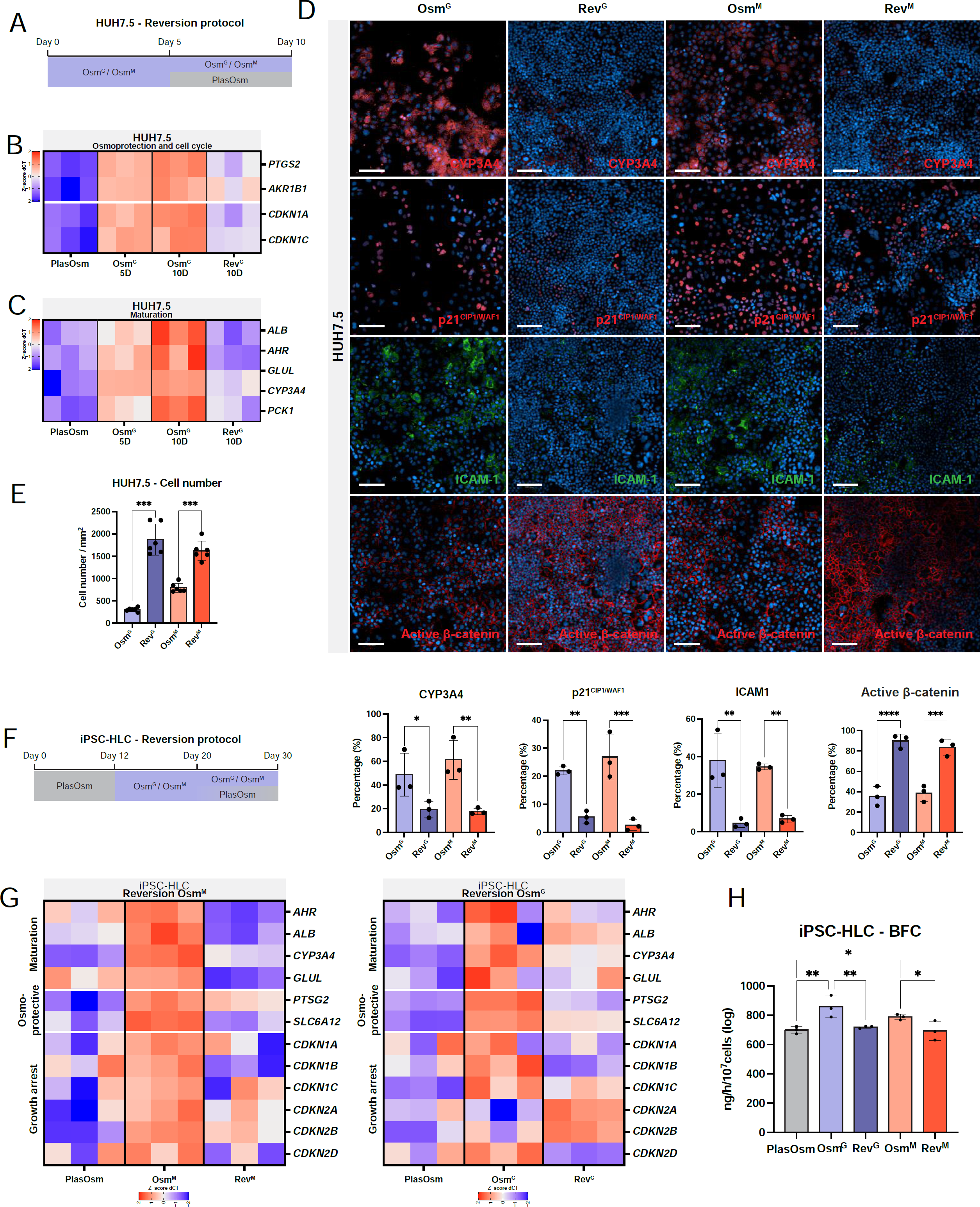
Prolonged hyperosmolarity is necessary for the stabilization of growth arrest and transcriptional maturation.

**(A)** Visual representation of HUH7.5 protocol of hyperosmolar withdrawal conditions. HypOsm media are swapped with PlasOsm conditions at day 5 and further cultured until day 10.
**(B)** Heatmap of osmoprotective genes and CDK inhibitors detected by RT-PCR at 5 and 10 days after Osm^G^ treatment and 5 days after Osm^G^ withdrawal represented in z-score of dCT values. N=3.
**(C)** Heatmap of hepatic maturation marker genes detected by RT-PCR at 5 and 10 days after Osm^G^ treatment and 5 days after Osm^G^ withdrawal represented in z-score of dCT values. N=3.
**(D)** Representative immunofluorescent images (up) and quantification of CYP3A4, p21^CIP1/WAF1^, ICAM-1 and active β-catenin (down) at day 10. HUH7.5 cells were cultured as indicated in (A). Statistics performed by Ordinary one-way ANOVA. N=3. Scale bar = 100 µm.
**(E)** Cell counts of HUH7.5 cells on day 10, 5 days of withdrawal of hyperosmolar conditions. Statistics performed by Brown-Forsythe and Welch ANOVA tests. N=6.
**(F)** Visual representation of hepatocyte-like cells differentiation protocol of hyperosmolar withdrawal conditions. At day 20, hyperosmolar conditions gradually reverted to PlasOsm levels until reaching day 30.
**(G)** Heatmap of maturation (upper), osmo-protective (middle) and growth arrest (bottom) genes detected by RT-PCR at D30 of hepatic differentiation treated with Osm^G^ (left) and Osm^M^ (right) for 18 days and after 10 days of hyperosmolar withdrawal, normalized to housekeeping gene *RPL19* and represented in z-score of dCT values. N=3.
**(H)** Functional comparison between D30 control HLCs, HLCs treated with Osm^G^ or Osm^M^ and HLCs after 10 days of Osm^G^ and Osm^M^ withdrawal for BFC metabolism. Statistics performed by Ordinary one-way ANOVA. N=3. All genes were normalized to housekeeping gene *RPL19*. All scale bars = 100 µm. All data represents mean ± SEM; *p < 0.05, **p < 0.01, ***p< 0.001, ****p<0.0001.

In line with the hepatoma line, transcriptional analysis of hiPSC-derived HLC differentiation demonstrated that the removal of hyperosmolar conditions from day 20 to day 30 (**Fig 6F**) of the differentiation protocol resulted in less quiescent and less mature progeny on day 30 (**Fig 6G**). Genes involved in growth arrest such as *CDKN1A*, *CDKN1B* and *CDKN1C* were downregulated upon relief of hyperosmolar stimulation (**Fig. 6G**). Furthermore, osmo-protective genes, such as *PTSG2* and *SLC6A12,* and hepatic maturation marker genes were downregulated suggesting that the osmotic protection was lost together with cell maturation in hiPSC-HLCs (**Fig. 6G**). Subsequently, this resulted in the loss of their detoxifying function picked up via CYP3A4-mediated BFC metabolism (**Fig. 6H**).

Therefore, upon removal from HypOsm conditions, both hepatoma HUH7.5 cells and hiPSC-HLCs regain a regular osmotic phenotype and lose growth arrest marker expression and the more mature phenotype achieved during the HypOsm treatment, indicating that cells in hyperosmolarity reside in a reversible quiescent state. This confirms that in order to achieve an improved hepatic phenotype and function following differentiation, HypOsm conditions need to be maintained over time to stabilize the more mature state and growth arrest.

### The adaptive osmo-protective response is a common mechanism shared by the mesodermal lineage

To validate whether the mechanism by which cells adopt an osmo-protective response to undergo cell cycle arrest is hepatocyte (endodermal lineage) specific, we further differentiated hiPSC to the mesodermal lineage, more specifically endothelial cells (ECs). Using a previously described protocol iETV2 hiPSC were differentiated to ECs^30^ (**Fig. S6A**). When cultured under HypOsm conditions (550 mOsm), no DNA damage occurred compared to control cultures, shown by the absence of pH2AX (**Fig. 7A**). Both Osm^G^ and Osm^M^ conditions induced a translocation of NFAT5 to the nucleus, as well as an increase in transcript levels for osmo-protective genes, such as *PTGS2* and *SCL6A12* (**Fig. 7B, 7C, S6B**). hiPSC-derived ECs cultured under HypOsm conditions proliferated less, visualized by a lowered cell count (**Fig. 7D**). This was in line with the decreased percentage of KI67^+^ cells in Osm^G^ conditions (**Fig. 7E**). In addition, endothelial cells in Osm^G^ expressed higher levels of several CDKi, including p15^INK4B^ (*CDKN2B*), p19^INK4D^ (*CDKN2D*), p21^CIP1/WAF1^ (*CDKN1A*) and p57^KIP2^ (*CDKN1C*) (**Fig. 7F**). We confirmed increased detection of p21^CIP1/WAF1^ positive cells at protein level in both Osm^G^ and Osm^M^ conditions (**Fig. 7G** and **7H**). Moreover, upon hyperosmolar treatment, endothelial-specific gene transcripts such as *KDR* and *LYVE-1* were notably higher expressed while there was a clear tendency of increased LYVE-1 protein levels (**Fig. 7I** and **7J**).

**Figure 7:**
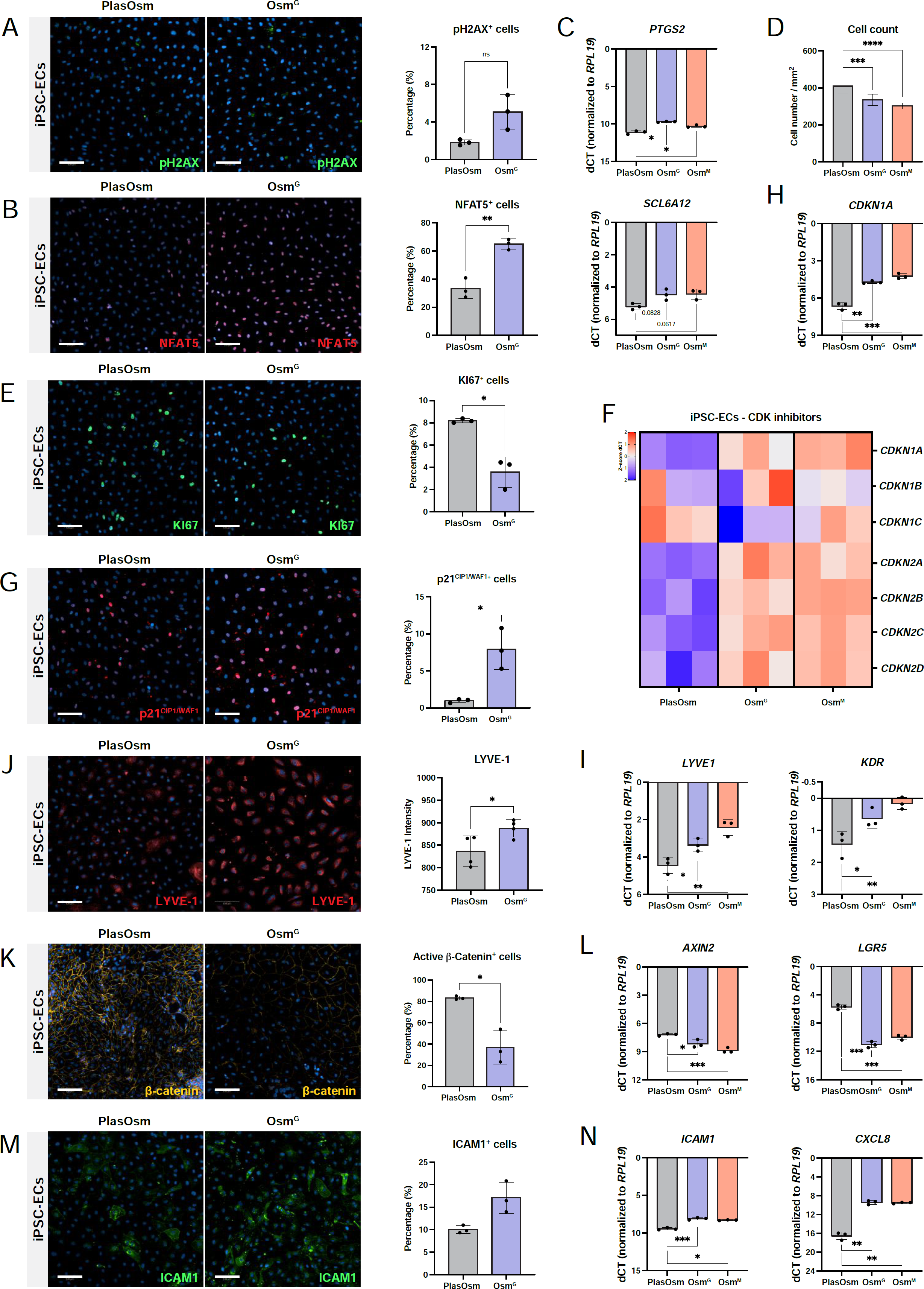
The adaptive osmo-protective response is a common mechanism shared by the mesodermal lineage.

**(A)** Representative immunofluorescent staining (left) and quantification (right) of pH2AX after 48 hours of Osm^G^ treatment in hiPSC-derived endothelial cells. Statistics performed by t-test with Welch’s correction. N=3.
**(B)** Representative immunofluorescent staining (left) and quantification (right) of NFAT5 of 48h Osm^G^ treatment in hiPSC-derived endothelial cells. Statistics performed by t-test with Welch’s correction. N=3.
**(C)** Gene expression of NFAT5 target genes 48h after Osm^G^ treatment detected by RT-PCR. Statistics performed by Brown-Forsythe and Welch ANOVA tests. N = 3.
**(D)** Cell count of cells treated with Osm^G^ and Osm^M^ after 48 hours. Statistics performed by Brown-Forsythe and Welch ANOVA tests. N = 6.
**(E)** Representative immunofluorescent staining (left) and quantification (right) of KI67 after 48 hours of Osm^G^ treatment in hiPSC-derived endothelial cells. Statistics performed by t-test with Welch’s correction. N=3.
**(F)** Heatmap of CDK inhibitors’ gene expression 10 days after Osm^G^ and Osm^M^ treatment detected by RT-PCR, normalized to housekeeping gene *RPL19*. Data visualized in z-score of dCT levels. N=3.
**(G)** Representative immunofluorescent staining (left) and quantification (right) of p21 after 48 hours of Osm^G^ treatment. Statistics performed by t-test with Welch’s correction. N=3.
**(H)** Gene expression of *CDKN1A* 48h after Osm^G^ treatment. Statistics performed by Brown-Forsythe and Welch ANOVA tests. N = 3.
**(I)** Representative immunofluorescent staining (left) and quantification (right) of LYVE1 detection by immunofluorescence of hiPSC-derived endothelial cells treated with Osm^G^ after 10 days in culture. Statistics performed by t-test. N=3.
**(J)** Gene expression of LYVE1 and KDR 48h after Osm^G^ treatment. Statistics performed by Brown-Forsythe and Welch ANOVA tests. N = 3.
**(K)** Representative immunofluorescent staining (left) and quantification (right) of active β-catenin after 48 hours of Osm^G^ treatment. Statistics performed by t-test with Welch’s correction. N=3.
**(L)** Gene expression of WNT target genes (*AXIN2* and *LGR5*) 48 hours after Osm^G^ treatment. Statistics performed by Brown-Forsythe and Welch ANOVA tests. N = 3.
**(M)** ICAM1 detection by immunofluorescence treated with Osm^G^ after 48 hours. Statistics performed by t-test with Welch’s correction. N=3.
**(N)** Gene expression of NF-κВ target genes (ICAM1 and CXCL8) 48 hours after Osm^G^ treatment. Statistics performed by Brown-Forsythe and Welch ANOVA tests. N = 3. All genes were normalized to housekeeping gene *RPL19*. All scale bars = 100 µm. All data represents mean ± SEM; *p < 0.05, **p < 0.01, ***p< 0.001, ****p<0.0001.

In line with the hepatocyte models, β-catenin levels together with WNT target genes *LGR5* and *AXIN2* were downregulated upon HypOsm conditions, displaying a suppressed WNT profile (**Fig. 7K** and **7L**). Additionally, the upregulation of *ICAM-1* and *CXCL8* indicated that Osm^G^ and Osm^M^ treatment of endothelial cells lead to the activation of NF-κВ signaling (**Fig. 7M, 7N,** and **S6C**). These results suggest that the cell cycle arrest and maturation in both HLC and endothelial differentiation are driven by a shared underlying mechanism.

Our results show that HypOsm culture conditions promote a cell cycle arrest and an enhanced transcriptional maturation during the differentiation of the two proposed differentiation models towards the endodermal and mesodermal lineage, suggesting that the HypOsm-induced cell cycle arrest might be used to improve the differentiation of hiPSC towards diverse lineages and cell types.

## DISCUSSION

As plasma osmolarity in vertebrates ranges from 285 to 300 mOsm, culture media commonly vary from 290 mOsm to 320 mOsm to prevent cells from undergoing apoptosis due to osmotic stress. However, cells in organs such as the liver, spleen, and kidneys reside in a physiological hyperosmolar range (**Fig. 1B**)^18, 19^. Here, we demonstrate that hepatoma cells and hiPSC undergoing differentiation to hepatocyte and endothelial-like cells can be cultured for prolonged periods in physiological HypOsm conditions without DNA damage or cell apoptosis, indicating that certain cell types can be *in vitro* cultured in HypOsm conditions mimicking the elevated osmolarity of wherein their *in vivo* counterparts reside. Moreover, our findings reveal that prolonged HypOsm culture conditions induce transcriptional and phenotypical changes resulting not only in the upregulation of osmolar responsive genes but also in growth arrest and subsequently in higher maturity which are two of the key functional characteristics of terminally differentiated cells.

Developing embryos are highly sensitive to osmotic changes ^70–72^. In agreement, we found that elevated hyperosmolarity (550 mOsm) induced cell death in pluripotent cells. However, these conditions did not affect the viability of hiPSC-derived hepatocytes or hepatoma cell lines. Our RNA-seq analysis comparing PSCs and PSC-derived hepatoblasts (day 12 differentiation protocol) showed that compared to undifferentiated hiPSC cells, hepatoblasts express higher levels of solute carrier (SLC) family genes, which are responsible for solute transport in the cells and are essential for adaptive osmolar responses ^73^. This indicates that the capacity to respond to hyperosmolar conditions is an intrinsic characteristic acquired during differentiation.

Multiple studies have shown a substantial amount of contradictory findings regarding the mechanisms underlying cellular response to osmolarity^17, 27, 51, 74, 75^. While some studies showed that HypOsm culture conditions induce DNA damage, p53 activation and cell death, others demonstrated that cells are able to adapt to HypOsm conditions by activating NFAT5, causing its nuclear localization where it drives the expression of osmo-protective genes. However, the majority of such studies only explored single HypOsm levels. We show that the resulting controversy can be explained by distinguishing between the effects caused by physiological versus toxic HypOsm levels. We first tested several HypOsm levels and confirmed that under toxic HypOsm conditions hepatoma cells increased p53 expression and underwent cell death. By contast, physiological HypOsm conditions did not prove to be harmful as it activated an osmolar transcriptional program mediated by an NFAT5 response and subsequently induced a protective cell cycle exit into the G_0_ phase where DNA damage, caspase activity or apoptosis could not be detected. As this was accompanied by a p53-independent increase in p21^CIP1/WAF1^ and CDKi levels, this indicated two completely opposite p53 responses under physiological and toxic HypOsm. Additionally, we found that physiological HypOsm culture conditions induced NF-κВ signaling and CDKi expression. When NF-κВ induction was inhibited, growth arrest did not occur and p21^CIP1/WAF1^ expression decreased, indicating that NF-κВ is a key player regulating HypOsm-induced growth arrest. Thus, in physiological hyperosmolar conditions, cell cycle exit caused by NF-κВ activation results in an adaptive cellular response, highlighting the importance of differentiating between physiological and toxic hyperosmolar conditions for future experimental approaches.

During development and adult tissue turnover, proliferation, and maturation are believed to occur concomitantly but diametrically opposed to one another^76–80^. While differentiating cells lose their self-renewable ability, proliferating tumorigenic cells undergo dedifferentiation^81–83^. Here, we found that the hyperosmotic growth arrest and the induced transcriptional maturation do not take place simultaneously, but rather that upregulation of mature genes is preceded by a cell cycle exit. Although we detected increased expression of multiple CDKi family member genes, p21^CIP1/WAF1^ was the most strongly upregulated cell cycle inhibitor in hepatoma cell lines and differentiation of hiPSC under hyperosmolar conditions. Previous reports have demonstrated that cell cycle exit is governed chiefly by the CDK inhibitor p21^CIP1/WAF1^ ^84, 85^. In adult mouse tissues, p21^CIP1/WAF1^ is found to be localized in terminally differentiated cells. Moreover, while p53-mediated p21^CIP1/WAF1^ expression is strongly linked to senescence, the p53-independent expression of p21^CIP1/WAF1^ has been correlated to guiding differentiation ^86–88^. In agreement with previous reports that NF-κВ regulates p21^CIP1/WAF1^ expression ^89^, we found that NF-κВ inhibition strongly reduced p21^CIP1/WAF1^ levels in hyperosmolar culture, resulting in accumulated cells in the S/G_2_ phases and a notable reduction in the transcriptional maturation. As the growth arrest mediated by p21^CIP1/WAF1^ and other CDKis preceded the expression of differentiation markers caused by hyperosmolar conditions, we suggest that enhanced differentiation might be a consequence of the preceding growth arrest. Although hPSC-HLCs showed a transcriptional cell cycle profile similar to plated PHH and increased maturation features, they did not yet fully resemble PHH at transcriptional maturation levels indicating that several factors are still lacking to further drive hPSC-HLCs towards their in vivo counterpart.

We previously demonstrated that amino acid supplementation improved HLC differentiation^29^. Here we show that the treatment of cells with the sugar-based osmolyte mannitol or multiple amino acids at high concentrations promotes an adaptive HypOsm response. We further demonstrated that both supplementations sequentially induced a growth arrest and cell maturation in hepatoma and hiPSC-HLCs. Furthermore, we disclose the molecular mechanism underlying the osmolarity-dependent growth arrest and improved maturation. Overall, we conclude that raising osmolarity levels under culture conditions, independently of the osmolyte, is the main force promoting growth arrest and cellular maturation from undifferentiated and pluripotent cells.

Comparison of the transcriptome of hiPSC-derived hepatoblasts with PHHs revealed lower transcriptional WNT levels in PHH, suggesting that WNT inhibition might promote cellular maturation from hepatoblast to cells with mature hepatocyte features. In line, Touboul et al. reported that inhibition of the WNT/β-catenin pathway is required to promote the differentiation of hepatoblasts into hepatocytes^90^. By contrast, activation of the canonical WNT/β-catenin pathway has been described for toxic HypOsm conditions causing apoptosis of mesangial cells^91^ or inner medullary collecting duct cells ^92^. Here, we found that physiological hyperosmolar conditions caused the repression of the WNT pathway at the protein and transcriptional level in hepatoma cells as well as HLCs and EC derived from hiPSC. Moreover, the osmolarity-dependent WNT repression was proven to be essential for hepatocyte transcriptional maturation, as reactivation of HypOsm-mediated WNT inhibition reduced the expression of hepatocyte markers. Surprisingly, although the WNT pathway plays a central role in regulating proliferation during development and in stem cell self-renewal, WNT reactivation did not affect cell proliferation or cell cycle markers, indicating that HypOsm-mediated WNT inhibition is necessary to promote transcriptional maturation but not growth arrest.

NF-kB activity levels have been reported to increase during differentiation ^93^ and mouse and human PSC missing key NF-κВ components cannot differentiate^94, 95^, emphasizing the importance of the NF-κВ pathway in mediating cellular differentiation. As NFAT5, induced under hyperosmolar conditions is a co-factor of the NF-κВ signaling cascade^96^, the inhibition of IKK2 suppresses the osmo-protective response as NFAT5 is unable to drive its target expression. Collectively, we show the inhibition of the NF-κВ pathway during HypOsm conditions prevents the cells from overcoming the hyperosmolar stimulus, disrupts cell cycle exit compromising their differentiation potential. Additionally, as NF-κВ inhibition reverted the cell cycle arrest in the hepatoma and hiPSC-HLCs, this possibly links the response to hyperosmolarity with the induced growth arrest. Although NF-κВ is commonly known to drive proliferation upon inflammation and tissue regeneration, several reports have proposed a potential crosstalk with p21^CIP1/WAF1^ regulating cell cycle exit^89, 97^. Moreover, NF-κВ has been reported to protect cells under irradiation via the upregulated p21^CIP1/WAF1^. Together with our findings, this suggests that NF-κВ-mediated protection towards hyperosmolarity is acquired throughout differentiation and contributes to a more homogeneous and improved differentiation outcome.

Differentiating patient-derived hiPSC to functional mature cell types such as liver cells is an attractive long-term aim in the regenerative medicine field and in the pharmaceutical industry for toxicology drug screenings. Hence, our findings have practical utility for more efficient generation of mature hiPSC-derived HLCs and ECs. Whether osmolarity might be used to modulate the generation of other mature cell types needs further investigation.

## ACKNOWLEDGMENTS

The authors would like to extend their gratitude to the Research Foundation – Flanders for the Ph.D. fellowships awarded to J.S.H.C. (1S65321N), PA (11M7822N), BV (11E7920N), FWO Research Project Grants G091521N, G073622N (FLL), FWO-SBO hiPSC-LIMIC S001121N (CV, FLL) and C1 KU Leuven internal grant C14/21/115 (FLL). We further thank Sebastian Munck and Axelle Kerstens from the Light Microscopy and Imaging Network LiMoNe, VIB Bio Imaging Core Leuven for the provided access, training, assistance and support to the use of their instruments and analysis software. We thank the laboratory of prof. Joris Vriens, and in particular Annelies Janssens for their assistance and use of their osmometer. Finally, we would like to thank Dr. Anna Osnato for her comments and critical reading of the manuscript.

## Supplementary Figure legends

**Supplementary Figure 1:**

**(A)** Gene expression of NFAT5 target genes detected by RT-PCR normalized to housekeeping gene *RPL19* across different levels of hyperosmolarity under Osm^G^. Statistical significance was determined compared to the control condition. Scale bar = 100 µm. N=3, statistics by Two-way ANOVA.
**(B)** Gene expression of osmo-adaptive genes detected by RT-PCR normalized to housekeeping gene RPL19 measured across different time points of hyperosmolarity (550 mOsm) under Osm^G^ and Osm^M^ treatment. Statistical significance was determined compared to the control condition. N=3, statistics by Brown-Forsythe and Welch ANOVA.
**(C)** Following cut-off values of FDR 0.05 and a fold change of 2, Differentially Expressed Genes were enriched for Gene Ontology terms after 36 hours of treatment. Full list of GO terms shown in (Supplementary Table 5).
**(D)** Following cut-off values of FDR 0.05 and a fold change of 2, Differentially Expressed Genes were enriched for Gene Ontology terms after 5 days of treatment. Full list of GO terms shown in (Supplementary Table 5).

**Supplementary Figure 2:**

**(A)** Visual representation of the FUCCI construct and scheme of cell cycle phases represented by the FUCCI colors. Adapted from Dolfi, et al. 2019^42^.
**(B)** Confocal images of FUCCI-HUH7.5 cells treated with increasing concentrations of indicated amino acids (up). Scale bar = 100 µm. Quantification of cell death (DAPI), cells in S/G2 phase, cells in G0/G1 phase and cells in cell cycle arrest represented in heatmaps (down).
**(C)** Western blot showing the protein levels of p53 and Actin following Osm^M^ treatment for 36 hours with increasing osmolarity levels.

**Supplementary Figure 3:**

**(A)** Boxplots representing the expression of the gene sets of specific gene ontology (GO) terms represented in the z-score of TPM values. Statistics by Mann-Whitney test. N = number of genes represented within each GO term and is indicated on represented boxplots. Full list of genes shown in Supplementary Table 6.
**(B)** Visual representation of the growth factor-based hepatic differentiation from induced PSCs^31^.
**(C)** Boxplots representing the expression of the gene sets of SCL gene ontology (GO) family represented in the z-score of TPM values. Statistics by Mann-Whitney test. N = 433 genes in the GO term. Full list of genes shown in (Supplementary Table 6).
**(D)** Gene Ontology (GO) enrichment of differentially expressed genes (DEGs) comparing PlasOsm vs HypOsm (Osm^G^) at D30 of differentiation. Full list of GO terms shown in Supplementary Table 8.
**(E)** Representative image (left) and quantification (right) of fluorescent detection of p53 and p21^CIP1/WAF1^ after 72h of Osm^G^ treatment. Scale bar = 100 µm. N=3, statistics by Two-Way ANOVA.
**(F)** Gene expression analysis of hepatic maturation markers upon Osm^M^ treatment in hiPSC-derived HLCs at D30 of differentiation. N = 3. Statistics by Welch’s t-test.
**(G)** Heatmap of hepatic maturation markers upon Osm^G^ treatment in hiPSC-derived HLCs at D30 of differentiation compared to PHHs plated.

All data represents mean ± SEM; *p < 0.05, **p < 0.01, ***p< 0.001, ****p<0.0001.

**Supplementary Figure 4: Wnt signaling regulates lineage specification during hyperosmolarity guided differentiation.**

**(A)** Gene ontology (GO) enrichment reveals differentially expressed signaling pathways between control and Osm^G^ treated cells at 36h after treatment. Full list of GO terms shown in Supplementary Table 9.
**(B)** Western blot of active and total β-catenin levels (upper panel) visualized and quantified to Actin levels (lower panel) after Osm^G^ or Osm^M^ treatment. N=3. Statistics by One-way ANOVA.
**(C)** Percentage of GFP+ HUH7.5 cells containing the WNT transcriptional reporter 7TGP treated with CHIR99021 in CTL and Osm^G^ conditions. N=3. Statistics by One-way ANOVA.
**(D)** Transcriptional levels of Wnt target genes across CTL, Osm^G^ and Osm^G^ + CHIR99021. Data is represented in z-score of TPM values.
**(E)** PCA plot of RNA-seq samples under CTL, Osm^G^ and Osm^G^ + CHIR99021 conditions.
**(F)** Heatmap of genes involved in maturation (left) and stemness/hepatocarcinoma (right) represented in z-score of TPM values across samples.
**(G)** Heatmap of genes involved in proliferation represented in z-score of TPM values across samples.
**(H)** Cell count of hepatoma HUH7.5 cell line at 0 and 36 hours and 5 days after CTL, Osm^G^ and Osm^G^ + CHIR99021 treatment. N=3. Statistics by Two-way ANOVA.
**(I)** KI67 detection by flow cytometry under Osm^G^ and Osm^G^ + CHIR99021 conditions after 36 hours. N=3, statistics by One-way ANOVA.
**(J)** Heatmap of genes involved in WNT signaling represented in z-score of TPM values across HLCs at D30 in PlasOsm conditions compared to PHHs plated.
**(K)** Gene expression analysis of hepatic maturation markers and WNT target genes after treatment with Osm^M^ and Osm^M^ + CHIR99021 at D30 of the differentiation protocol. Data is represented in z-score of the dCT levels. N=3.
**(L)** Gene expression analysisof genes related to proliferation after treatment with Osm^M^ and Osm^M^ + CHIR99021 at D30 of the differentiation protocol. N=3. Statistics by Welch’s t-test.

All data represents mean ± SEM; *p < 0.05, **p < 0.01, ***p< 0.001, ****p<0.0001.

**Supplementary Figure 5:**

**(A)** Transcription factor prediction using the TTRUST tool on differentially expressed genes at 36 hours of glycine treatment.
**(B)** Heatmaps of bulk RNA-seq data representing the z-score of TPM values across presented sample conditions showing genes associated with NF-κВ signaling comparing hiPSC-HLCs under PlasOsm at day 30 to PHHs.
**(C)** Representative images (up) and quantification (down) of immunofluorescent detection of IκВα and ICAM-1 under PlasOsm, Osm^G^ and Osm^M^ conditions in HUH7.5 cells. Scale bar = 100 µm. N=3, statistics by Brown-Forsythe and Welch ANOVA tests.
**(D)** Heatmaps of bulk RNA-seq data representing the z-score of TPM values across presented sample conditions showing genes associated with NF-κВ signaling comparing hiPSC-HLCs under PlasOsm and Osm^G^ treatment.
**(E)** Representative images (left) and quantification (right) of immunofluorescent staining of p21^CIP1/WAF1^ under PlasOsm, Osm^G^ and Osm^G^ + TPCA conditions, as indicated in HUH7.5 cells. Scale bar = 100 µm. N=3, statistics by Brown-Forsythe and Welch ANOVA tests.
**(F)** Principal component analysis of bulk RNA-seq data of PlasOsm, Osm^G^ and Osm^G^ + TPCA treated HUH7.5 cells after 36 hours of treatment.
**(G)** Heatmaps of bulk RNA-seq data representing the z-score of TPM values across presented sample conditions of HUH7.5 cells showing genes associated with NF-κВ signaling and osmolar response.
**(H)** Heatmaps of bulk RNA-seq data representing the z-score of TPM values across presented sample conditions of HUH7.5 cells showing genes associated with ER stress.
**(I)** Gene ontology (GO) enrichment of upregulated DEGs in TPCA-1 treated HUH7.5 cells under Osm^G^ conditions. Full list of GO terms shown in (Supplementary Table 14).

All data represents mean ± SEM; *p < 0.05, **p < 0.01, ***p< 0.001, ****p<0.0001.

**Supplementary Figure 6: The adaptive osmo-protective response is a common mechanism shared by the mesodermal lineage.**

**(A)** Visual representation of the growth factor-based endothelial differentiation from pluripotent hiPSC.
**(B)** NFAT5 detection by immunofluorescence treated with Osm^M^ after 48 hours. Statistics performed by t-test with Welch’s correction. N=3.
**(C)** ICAM1 detection by immunofluorescence treated with Osm^M^ after 48 hours. Statistics performed by t-test with Welch’s correction. N=3.

All genes were normalized to housekeeping gene *RPL19*. All scale bars = 100 µm. All data represents mean ± SEM; *p < 0.05, **p < 0.01, ***p< 0.001, ****p<0.0001. Controls depicted in this supplementary figure are the same as in Figure 7.

## Supplementary Tables

**Table 1** - Antibody list.

**Table 2** - Primer list.

**Table 3** - Defined gene sets used in heatmaps.

**Table 4** - DEGs HUH7.5 PlasOsm vs OsmG.

**Table 5** - GO terms HUH7.5 PlasmOsm vs OsmG.

**Table 6** - Gene lists defined by GO terms used in heatmaps.

**Table 7** - DEGs HLC PlasmOsm vs OsmG.

**Table 8** - GO terms HLC PlasmOsm vs OsmG.

**Table 9** - Identified signaling pathways based on DEGs HUH7.5 36h.

**Table 10** - PC axes HUH7.5 OsmG vs OsmG + CHIR.

**Table 11** - TTRUST enrichment HUH7.5 OsmG 36h.

**Table 12** - DEGs HUH7.5 OsmG vs OsmG + TPCA36h.

**Table 13** - DEGs HLC OsmG vs OsmG + TPCA 72h.

**Table 13** - GO terms HUH7.5 OsmG vs OsmG + TPCA 36h.

