## Supplementary figures and images for "Osmolar modulation drives reversible cell cycle exit and human pluripotent cell differentiation via NF-κВ and WNT signaling"

### Supplementary Figure 1

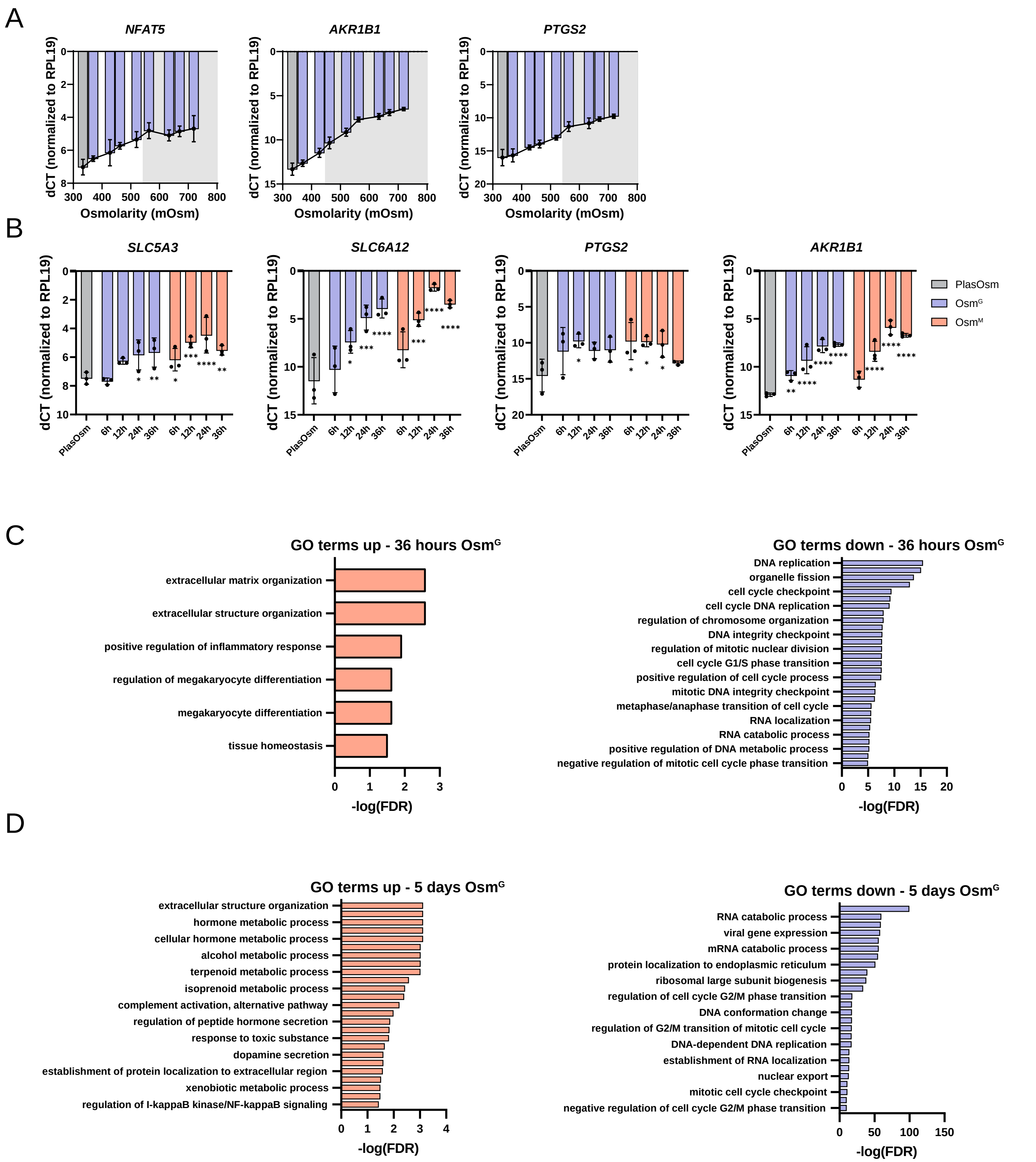

### Supplementary Figure 2

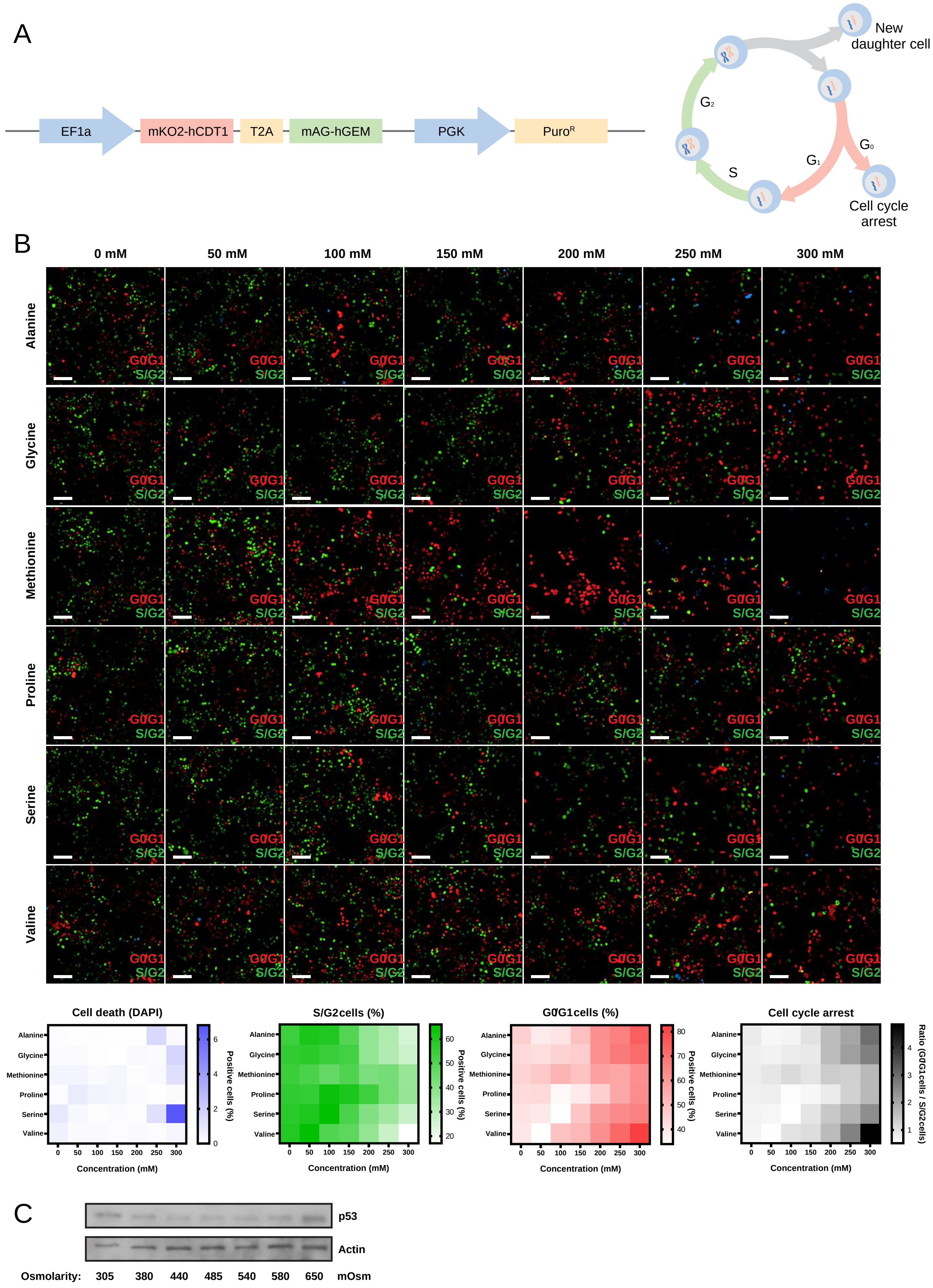

### Supplementary Figure 3

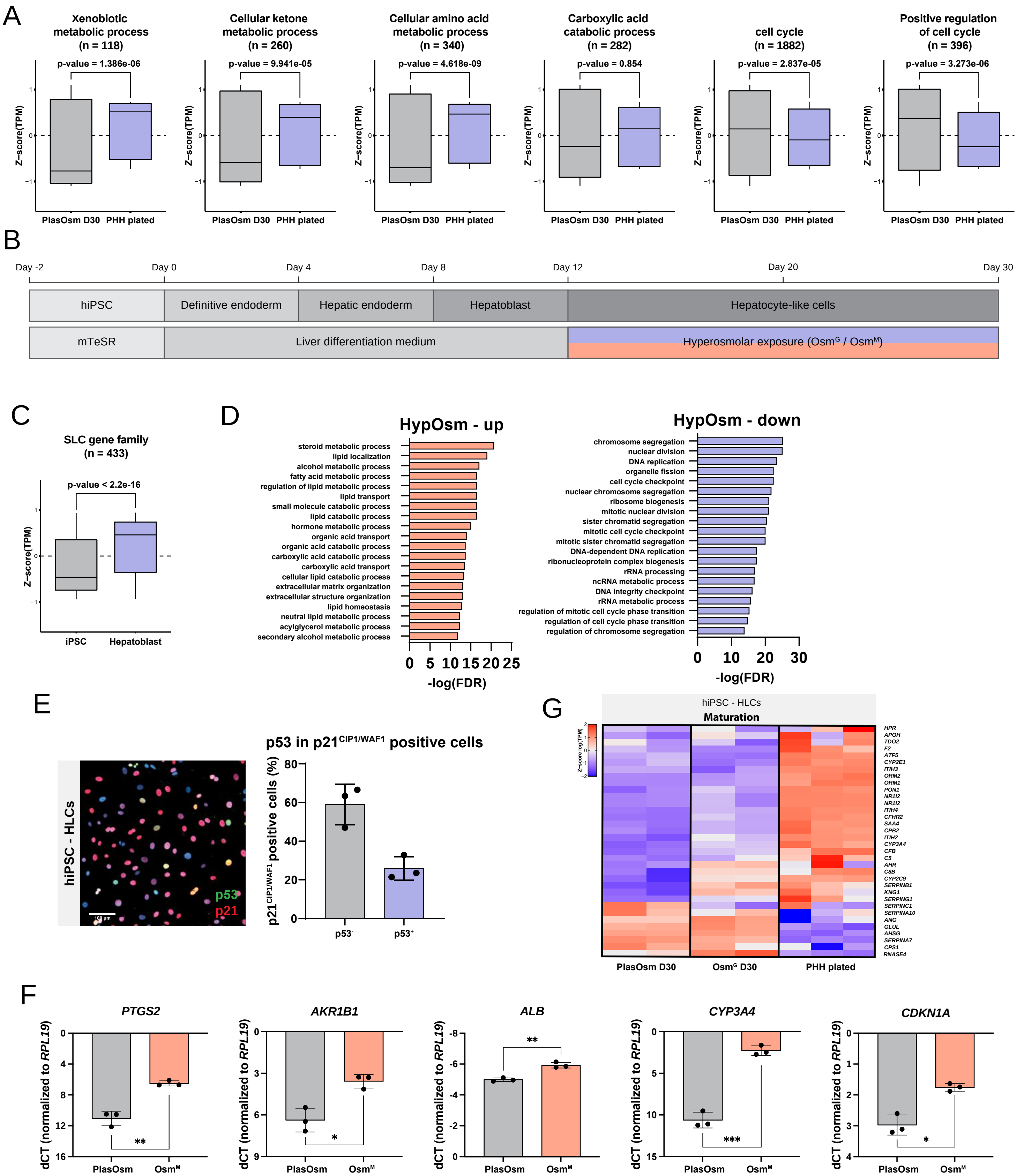

### Supplementary Figure 4

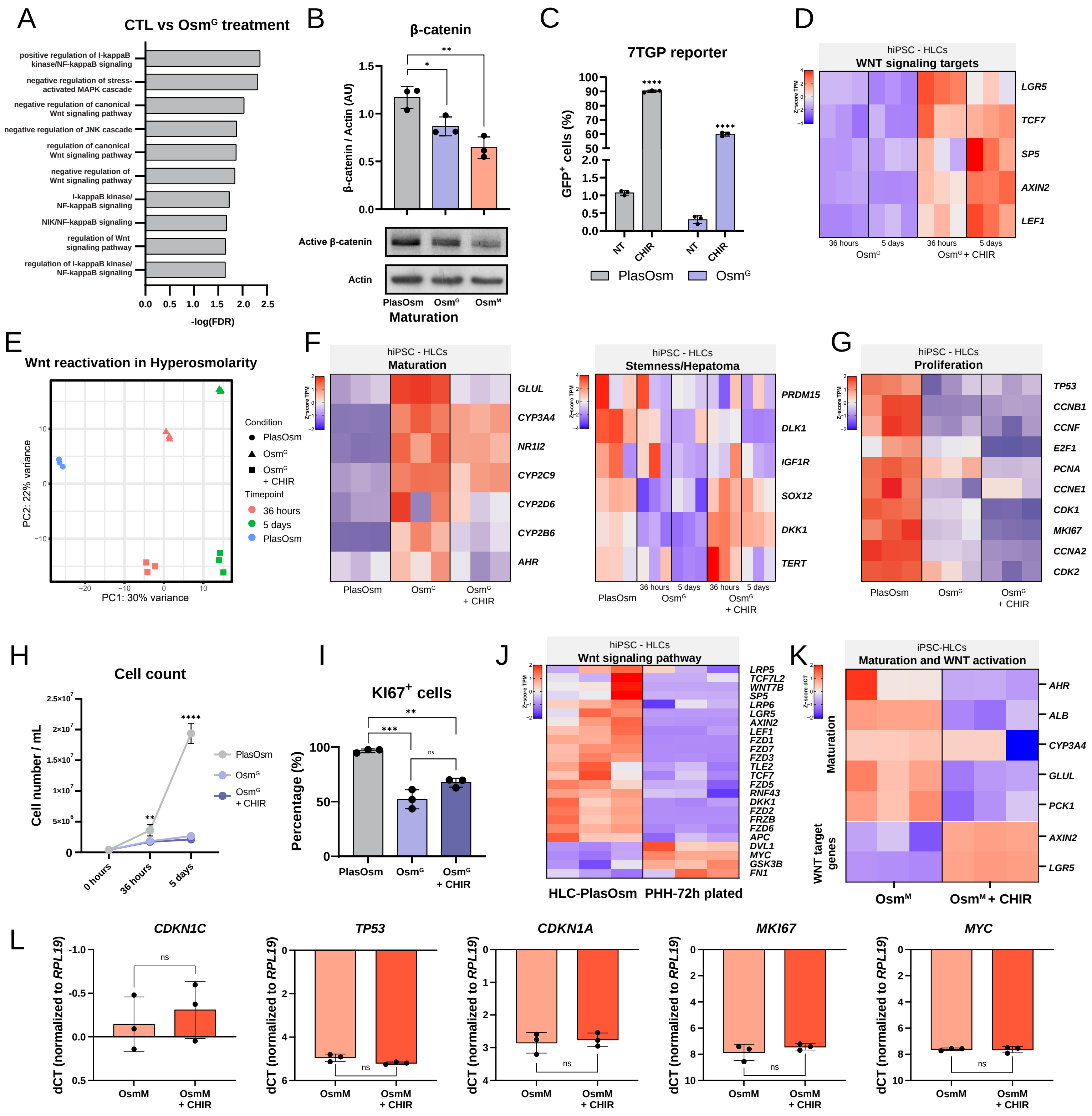

### Supplementary Figure 5

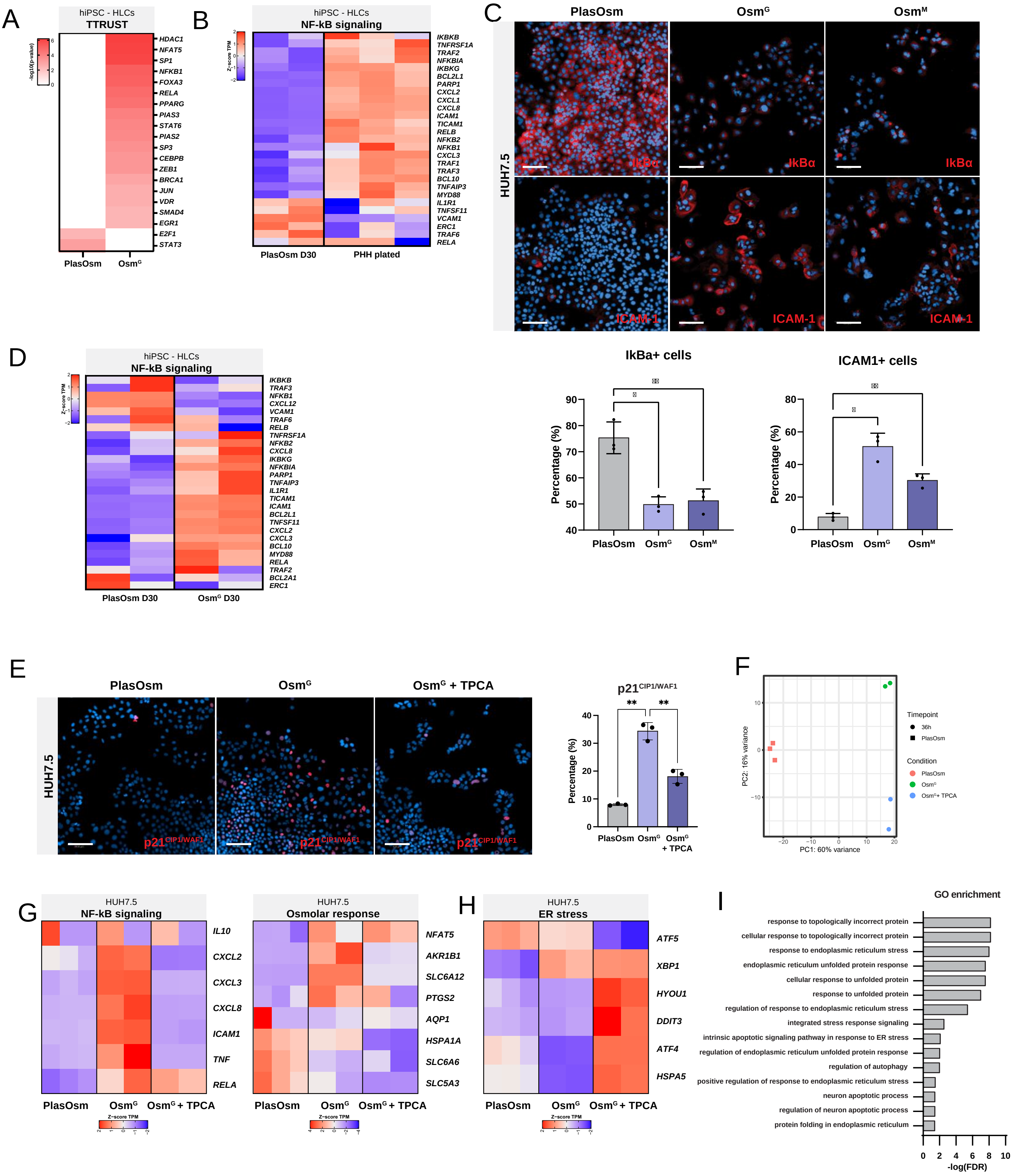

### Supplementary Figure 6

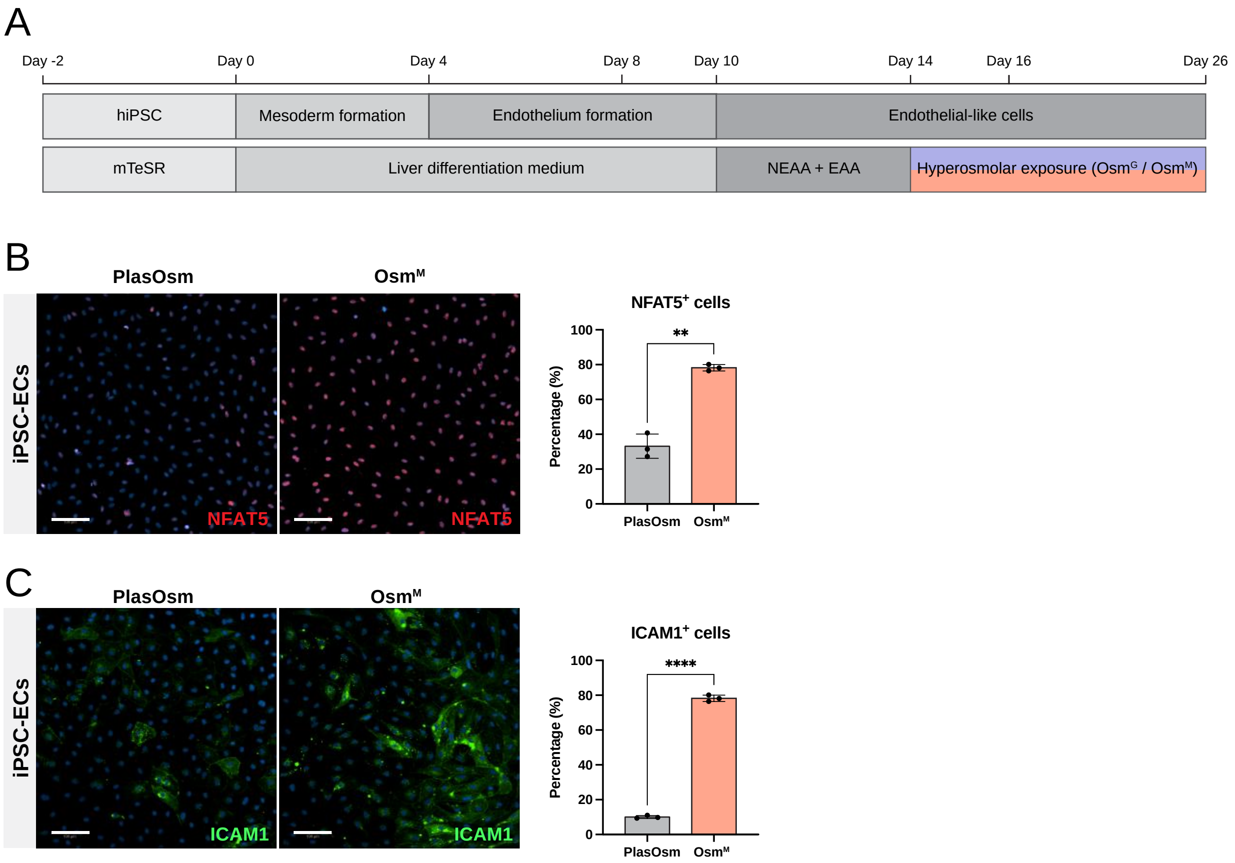
